# Phylogenomics resolves a 100-year-old debate regarding the evolutionary history of caddisflies (Insecta: Trichoptera)

**DOI:** 10.1101/2024.02.26.582007

**Authors:** Xinyu Ge, Lang Peng, John C. Morse, Jingyuan Wang, Haoming Zang, Lianfang Yang, Changhai Sun, Beixin Wang

## Abstract

Trichoptera (caddisfly) phylogeny provides an interesting example of aquatic insect evolution, with rich ecological diversification, especially for underwater architecture. Trichoptera provide numerous critical ecosystem services and are also one of the most important groups of aquatic insects for assessing water quality. The phylogenetic relationships of Trichoptera have been debated for nearly a century. In particular, the phylogenetic position of the “cocoon-makers” within Trichoptera has long been contested. Here, we designed a universal single-copy orthologue and sets of ultraconserved element markers specific for Trichoptera for the first time. Simultaneously, we reconstructed the phylogenetic relationship of Trichoptera based on genome data from 111 species, representing 29 families and 71 genera. Our phylogenetic inference clarifies the probable phylogenetic relationships of “cocoon-makers” within Integripalpia. Hydroptilidae is considered as the basal lineage within Integripalpia, and the families Glossosomatidae, Hydrobiosidae, and Rhyacophilidae formed a monophyletic clade, sister to the integripalpian subterorder Phryganides. The resulting divergence time and ancestral state reconstruction suggest that the most recent common ancestor of Trichoptera appeared in the early Permian and that diversification was strongly correlated with habitat change.

## 1. Introduction

Aquatic insects are an example of the evolutionary process from the aquatic (marine) to terrestrial to aquatic (freshwater) environments, and account for 60% of known freshwater species (Dijkstra et al., 2014; Li et al., 2001; Morse, 2017). In order to adapt to the freshwater environment, aquatic insects have evolved a variety of specialized body structures, physiological processes, behaviors, including swimming legs and body movements, respiratory tubes tracheal gills, osmoregulatory functions, and habitat-specific behaviors, among which Trichoptera provide an excellent example of diversity in aquatic insect evolution (Holzenthal et al., 2011; Morse et al., 2019; Wiggins, 1996). Trichoptera inhabit a wide variety of aquatic environments, including freshwater streams, rivers and lakes, wetlands, intertidal zones, and marine tidal pools. To survive in diverse environments, Trichoptera are as varied in their nutrient sources and feeding behaviors as freshwater Diptera and employ silk for habitat modification in more ways than any other group of animals, for example prompting their popular recognition as underwater architects (Anderson and Sedell, 1979; Mackay and Wiggins, 1979; Wiggins, 2004; Fig. 1). Over the course of at least 260 million years of evolution, they have exhibited extraordinary morphological, taxonomic, and ecological diversity, with about 52 families and more than 17,000 species described worldwide (Morse, 2023). Most Trichoptera have aquatic egg, larval, and pupal life stages, and terrestrial adults, each with highly differentiated morphological structures reflecting their niche adaptations (Morse et al., 2019).

**Fig. 1.**
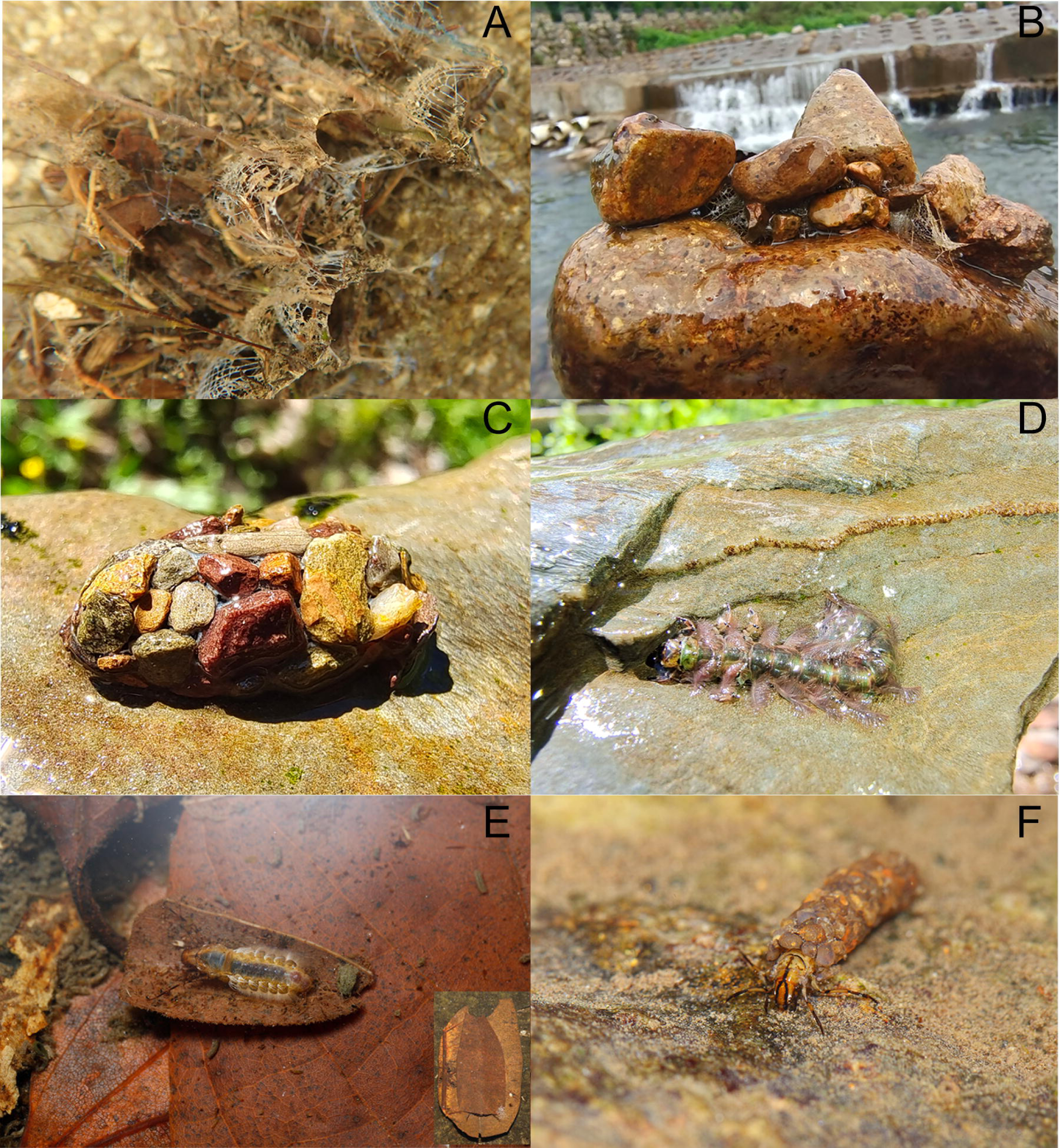
The retreat and case of caddisflies. A: retreat of *Arctopsyche* sp. (Hydropsychidae), from Zhejiang, China; B: retreat of *Stenopsyche* sp., (Stenopsychidae), from Zhejiang, China; C: saddle-case of *Glossosoma* sp., (Glossosomatidae), from Zhejiang, China; D: free-living *Himalopsyche malenanda*, (Rhyacophilidae), from Qinghai, China; E: tubular case of *Anisocentropus* sp., (Calamoceratidae), from Guangdong, China; F: tubular case of *Psilotreta* sp., (Odontoceridae), from Guangdong, China; A-D photograph by Haoming Zang; E, F photograph by Qianle Lu.

A hundred years ago, Trichoptera were divided into two suborders (Annulipalpia and Integripalpia), based on the presence or absence of an annulate apical segment of each maxillary and labial palp, a classification which is still valid today (Martynov, 1924; Morse, 1997). However, there are some special groups, now known as “cocoon-makers”, including families Glossosomatidae, Hydrobiosidae, Hydroptilidae, Rhyacophilidae, and Ptilocolepidae whose phylogenetic relationships have puzzled trichopteran scholars for many years. At first, these families were thought to belong to the Integripalpia (Morse, 1997; Ross, 1956). Subsequently, due to the hypothesis that semi-permeable pupal cocoons were considered primitive, “cocoon-makers” were regarded as the earliest grade in Trichoptera (Weaver, 1984; Wiggins and Wichard, 1989). Another hypothesis based on the semi-permeable cocoon hypothesis and caddisfly species morphology divided them, placing some at the bases of each of the two suborders (Frania and Wiggins, 1997). Early studies based on a few DNA markers showed that “cocoon-makers” were considered as a basal grade of Integripalpia that was sister to the monophyletic integripalpian subterorder Phryganides (with larvae generally constructing portable tubular cases); these studies considered sister families Hydroptilidae and Ptilocolepidae to be the earliest lineage in Integripalpia (Kjer et al., 2016; Thomas et al., 2020). Some more-recent studies based on the mitogenome and five molecular markers (*COI*, *18S rRNA*, and three nuclear genes) suggested that Hydroptilidae separated from other “cocoon-makers” and formed a sister group with Annulipalpia, contrary to the findings of Kjer et al (2016) and Thomas et al (2020) (Ge et al., 2023; Grigoropoulou et al., 2022). In addition, the phylogenetic positions of the families Glossosomatidae, Hydrobiosidae, and Rhyacophilidae as well as their relationships with each other and with Phryganides have not been determined, whether based on a few molecular markers or on the mitochondrial genomes (Ge et al., 2023; Thomas et al., 2020). To date, most molecular phylogenetic studies of Trichoptera have sampled only 15 or fewer genes, based on a matrix of less than 15,000 nucleotides in length. Although this limited genetic sampling has helped to reveal or support relationships between the suborders, among some superfamilies, and families, it has been insufficient to address with strong support some of the deep evolutionary relationships within Trichoptera, such as the phylogenic relationships within Psychomyioidea (Chamorro and Holzenthal, 2011; Johanson and Espeland, 2010; Johanson et al., 2012; Kjer et al., 2016; Thomas et al., 2020). The date of origin of the order is also uncertain due to unstable or faulty phylogenetic relationships (Malm et al., 2013; Thomas et al., 2020, 2023). Therefore, resolving the phylogenetic relationships and timescale of Trichoptera evolution has proven challenging, and revealing phylogenetic relationships of these lineages is critical to understanding early trichopteran character evolution, and aquatic environment adaptation.

In recent years, the development of phylogenomics has brought us new hope to solve the phylogenetic relationship of Trichoptera. In contrast to the problems that small molecular markers and mitochondrial genomic markers may have in reconstructing ancient nodes, obtaining thousands of loci through phylogenomics methods, sampling Universal Single-copy Orthologs (USCOs; Yu et al., 2022; Zhang et al., 2019), Anchored Hybrid Enrichment (AHE; Branstetter et al., 2017; Faircloth et al., 2012; Lemmon et al., 2012), and Ultraconserved Elements (UCEs; Zhang et al., 2023) can help overcome recalcitrant nodes on the tree of life (Kumar et al., 2012; Williams et al., 2020; Young and Gillung, 2020). In fact, phylogenomics has also greatly improved our understanding of the origin and evolution of insects (Misof et al., 2014). Many scholars have developed different USCO datasets, AHE probes, and UCE probes by combining different research groups (Diptera, Hemiptera, Coleoptera, Hymenoptera, Collembola; Branstetter et al., 2017; Faircloth, 2017; Godeiro et al., 2023; Sun et al., 2020; Waterhouse et al., 2018). In the transcriptome studies being conducted at the same time as ours, AHE has been used to reconstruct phylogenetic relationships of Trichoptera (Frandsen et al., 2023).

Therefore, to resolve the controversy about the phylogenetic relationships of the cocoon-maker families in Trichoptera, this study reconstructed the phylogenetic relationship of Trichoptera based on phylogenomics. The first Trichoptera USCO dataset and UCE probe sets were developed using high-quality assembly and protein reference genes. Herein, we report results of our analysis of newly sequenced low-coverage whole-genome data for 86 trichopteran species, having extracted thousands of USCOs and UCEs from whole-genome data to compile the largest genomic dataset of Trichoptera compiled to date. To reach this milestone, we filtered the loci using different strategies and different models to build phylogenetic trees with reduced probability of systematic errors. Concurrently, we discuss the validity of these two markers, propose a new hypothesis, and elucidate reliable phylogenetic relationships. Based on the new phylogenetic framework of the system and the fossil evidence, the divergence time was inferred, and the evolution of the key shape features was hypothesized.

## 2. Materials and methods

### 2.1 Taxon sampling and Molecular techniques

We collected 86 trichopteran species from 61 genera in 28 families, employing ultraviolet traps, and Malaise traps for adults and D-frame aquatic nets for larvae during 2017 to 2022 (Table S1). Specimen identification were conducted by Xinyu Ge, Lang Peng and Changhai Sun. We also downloaded 25 trichopteran genomes from GeneBank (as of 1 June 2023), for a total of 111 Trichoptera genomes as ingroups (Table S2). According to the phylogeny of Lepidoptera, the genomes of four families (Micropterigidae, Tineidae, Psychidae, and Choreutidae) were selected as outgroups for phylogenetic reconstruction (Kawahara et al., 2019). The newly sequenced samples were subjected to DNA extraction using the DNeasy Blood and Tissue kit (QIAGEN). All voucher specimens are stored in the Insect Museum of Nanjing Agricultural University, Nanjing, Jiangsu Province, China. Sequencing libraries were prepared using a library preparation kit and separate Illumina and BGI sequencing libraries were prepared for each sample according to the vendor’s protocols. We performed paired-end 150 bp sequencing for each library, with an insertion fragment length of 350 bp. Given the substantial variations in genome size among Trichoptera, we systematically screened the 28 collected families. We prioritized families lacking published genomes or those with inadequately represented genome species. Representative species within these families were selected for genome size assessment; the sequencing volume for these representative species ranged from 30 to 80 Gb. Following a comprehensive statistical analysis of the outcomes, it was observed that each library of the remaining species yielded approximately 10– 50 Gb of raw data, ensuring a sequencing coverage of more than 20×.

### 2.2 Genome size evaluation and assembly

Genome size assessment was conducted based on the frequency distribution of k-mer. Initially, BBmap v38.67 (Bushnell, 2014) was employed to eliminate repetitive sequences (clumpify.sh) and remove low-quality sequences (bbduk.sh). Subsequently, k-mer distribution values were computed using khist.sh with the parameter k=21. The final assessment of the genome involved using the R package within GenomeScope v2.0.1 (Vurture et al., 2017) to calculate k-mer distribution and heterozygosity, with the maximum sequencing coverage set at 10,000. PLWS v1.0.7 was used to assemble trichopteran genomes (Zhang et al., 2019). Firstly, the raw data underwent quality control using the aforementioned methods. Whereafter, the genome was assembled using Minia v3.2.4 (Chikhi and Rizk, 2013) with k-mer values ranging from 21 to 121. Redundans v0.13c (Pryszcz and Gabaldón, 2016) was used to remove redundant contigs. Ultimately, BESST v2.2.8 (Sahlin et al., 2014) and GapCloser v1.12 (Luo et al., 2012) were employed for extension and gap filling of sequence, respectively. The newly assembly trichopteran genome have been deposited on the National Genomics Data Center (NGDC).

### 2.3 USCO dataset and UCE probe design

The development of the trichopteran USCOs dataset was conducted following the design workflow published by Sun et al. (2020). The protein sequences of 10 trichopteran species and 4 lepidopteran species were downloaded from Gigabase and GeneBank (Table S3). The completeness of the downloaded protein sequences was evaluated using BUSCO v3.0.2 (Waterhouse et al., 2018) in protein mode (-m protein). Protein clustering was executed utilizing OrthoFinder v2.3.8 (Emms and Kelly, 2019), Subsequent procedures encompassed alignment, trimming, modeling of conserved regions, and sequence annotation, collectively leading to the establishment of the trichopteran USCO dataset (Trichoptera_odb1). The development of the UCE probe set followed the workflow by Faircloth (2017) and Zhang et al. (2019). Genomes from 17 trichopteran species and one lepidopteran species were selected for UCE probe design, as detailed in Table S4. The highest-quality genome (*Limnephilus lunatus* Curtis; Limnephilidae) was selected as the base genome for alignment. Subsequently, genomes were resampled, and the base genomes were aligned using ART-20160605 (Huang et al., 2012) and Stampy v1.0.32 (Lunter and Goodson, 2011), respectively. Ultimately, the UCE probes were designed using PHYLUCE v1.6.6 (Faircloth, 2017). The final baits at conserved sites must be shared by at least 15 species to ensure their suitability for subsequent analyses.

### 2.4 USCO, UCE extraction extract and Matrix preparation

BUSCO was employed to extract USCOs for all taxa, using the newly generated Trichoptera_odb1 dataset (n = 3860) with the parameter “-m genome”. To obtain more “complete” loci, the standard deviation for “lengths_cutoff” was increased by one-fold. As a preliminary filtering step for the loci, those with fewer than two sequences were excluded. MAGUS v0.1.1 (Smirnov and Warnow, 2020) was used with MAFFT to conduct homologous region alignment of the amino acid sequences of USCO. The ClipKit v1.1.5with the kpi strategy was then applied to retain parsimony-informative sites from the results of alignment.

To reduce systematic errors, two strategies were sequentially employed for gene filtering in this study. The first strategy was applied for filtering based on the gene characteristic, involving the following steps: (1) Phykit v1.2.1 (Steenwyk et al., 2021) was used to detect the number of concise information sites for each locus, and loci with more than 100 concise information sites were retained; (2) filtering was based on the homogeneity of each sequence: Phykit was used to assess the relative composition variability (RCV) values; (3) IQ-TREE v2.2.2.7 (Minh et al., 2020) was employed to exclude sequences that deviated from the assumptions of stationarity, reversibility, and homogeneity (SRH) of the loci. The parameter “—symtest” was used, and loci were retained based on an intermediate p-value of 0.05. Subsequently, Phykit was used to generate taxon-occupancy matrices at 60%, 70%, and 80%, denoted as USCO60/70/80. The second filtering strategy involved filtering based on the gene tree features for each gene: (1) The phylogenetic tree of each gene was constructed using IQ-TREE, setting the EX_EHO model and 1000 replicates of UFBoot2 (Hoang et al., 2017) applied to the USCO60/70/80 matrices; (2) Treeshrink v1.3.7 (Mai and Mirarab, 2018) was employed to identify and remove abnormally long branches in each gene tree, indicative of potentially paralogous sequences and assembly errors, with the parameter “-q 0.05”. After obtaining the new gene sequences, manual verification and the reconstruction of the phylogenetic tree for each gene, based on the EX_EHO model, were performed; (3) genes with an average bootstrap support (ABS) value greater than 75 were retained, and datasets USCO60/70/80_abs75 were generated for subsequent filtering. (4) The Degree of Violation of the Molecular Clock (DVMC) based on the molecular clock hypothesis, Phykit was employed for filtering based on the Degree of Violation of the Molecular Clock (DVMC) and treeness (proportion of the tree distance found on internal branches). Finally, FASconCAT-G v1.04 (Kück et al., 2014) was used to concatenate the retained loci, facilitating subsequent phylogenetic analysis.

A custom script developed by Zhang et al., (2019) was employed to extract UCEs for each species. The initial input files encompassed all 115 assemblies along with the newly generated trichopteran UCE probe set. The filtering strategy for UCE closely resembled that applied to USCO. UCEs with fewer than two sequences were excluded, and MAFFT was employed for sequence alignment using the L-INS-I strategy. The kpi strategy in ClipKit was used to retain parsimony-informative sites. Phykit was used to retain loci with a count of concise information sites greater than 100, and RCV heterogeneity tests were employed for further filtering. The SRH test with a cutoff parameter of 0.05 was performed using IQ-TREE. Taxon-occupancy matrices at 50%, 70%, and 90% were generated. IQ-TREE was employed to infer individual gene trees using a GTR model, and the Treeshrink was used to scrutinize long branches. Loci with ABS values greater than 70 were selected through subsequent analyses, resulting in datasets labeled UCE50/70/90_abs70. The DVMC and treeness tests were then applied for final filtration. In the final step, the loci were concatenated using FASconCAT-G for each matrix.

### 2.5 Phylogenetic analyses

For the data matrices obtained from two different types of molecular markers, USCO and UCE, we employed a series of models and various calculation methods to address issues such as rate heterogeneity, lineage heterogeneity, and incomplete lineage sorting (ILS) that could potentially affect phylogenetic reconstruction. To address the systematic errors caused by incomplete lineage sorting, we used the Multi-species coalescent model (MSCM). All gene trees within each data matrix of the two molecular markers were inputted into ASTER v1.15, employing w-astral strategy (Zhang and Mirarab 2022). This approach was employed to infer species trees for different matrices and estimate branch support rates.

The IQ-TREE was used to infer Maximum Likelihood (ML) trees for both the USCO and UCE matrices. The best-fitting substitution models for each gene partition were evaluated using the MODELFINDER module (Kalyaanamoorthy et al., 2017) integrated into IQ-TREE. The best model for each partition was determined, constrained to the specified LG for USCO matrices and GTR for UCE matrices, with the relaxed algorithm “-rclusterf 10”. Then, the GHOST (General Heterogeneous evolution On a Single Topology; Crotty et al., 2019) model was applied with “-m LG+FO+H4 and GTR+FO+H4” for the USCO and the UCE matrices, respectively. To alleviate the impact of data heterogeneity on phylogenetic reconstruction results, the EX_EHO mixture model (EX_EHO+FO+R) and the PMSF (Posterior Mean Site Frequency; Wang et al., 2017) model were employed for each USCO matrix in IQ-TREE. In the tree-building process based on the PMSF model, to ensure accuracy and eliminate the impact of guide trees with different topologies on PMSF tree inference, ML trees constructed using the partitioning model, the EX_EHO mixture model from the USCOs, and the ML tree of the partitioning model of UCEs were used as initial guide trees. The resulting tree was then used as the new guide tree for a second round of PMSF phylogenetic tree inference, repeating the process at least two times. To further reduce influence of multiple substitutions saturation, the Dayhoff6 recoding strategy was applied to each USCO matrix. Phylogears v2.2.0 (Tanabe, 2008) was used to convert 20 amino acids into 6 coding states (0–5), and the recoded sequences were then imported into IQ-TREE for analysis, with the model parameters set as “-m GTR+R”. Bayesian inference (BI) was conducted using PhyloBayes MPI v1.8c (Lartillot et al., 2013). The CAT+GTR model was employed based on USCO80_abs75, and two Markov chain Monte Carlo chains were run until achieving an effective size (>50) and convergence (maxdiff < 0.1). Finally, a strict consensus tree was generated after discarding the initial 25% of trees as burn-in.

The genealogical concordance was calculated using the gCF (gene concordance factor) and the sCF (site concordance factor) given the reference tree and gene trees using IQ-TREE. Additionally, to verify the reliability of different topologies generated by ML analysis in this study, the Approximately Unbiased test, weighted Kishino–Hasegawa test, and weighted Shimodaira– Hasegawa test were performed using in IQ-TREE (Kishino and Hasegawa, 1989; Shimodaira and Hasegawa, 1999; Shimodaira, 2002). The USCO70_abs75 matrix and the PMSF model (-m LG+C60+F+G) were chosen for this analysis, with parameters set as “-zb 1000 -zw -au”.

### 2.6 Divergence time estimation

The divergence time estimation used the PAML v4.9j plugin MCMCTREE (Yang, 2007). The PMSF tree generated from the USCO70_abs75 matrix served as the input topology for this analysis. Fossil calibration points were selected by searching the Paleobiology Database (PBDB; https://paleobiodb.org/navigator/). 12 fossil calibration points were marked in the input tree file: A Late Carboniferous fossil served as the root calibration point with a calibrated time of 322 million years ago (Ma). The oldest fossil of Trichoptera from the Late Triassic, Terrindusia sp., was set as the calibration point for the common ancestor of Trichoptera (Zheng et al., 2018). The calibration point for the common ancestor of Trichoptera was set with a broad range of 237–314 Ma. A fossil from the Early Jurassic Hettangian stage, representing the Glossata suborder of Lepidoptera, was set as the calibration point for the common ancestor of Lepidoptera with a range of 314–201 Ma (van Eldijk et al., 2018). Other fossil correction points and reference fossils are detailed Table S5. Hessian matrices were quantified using the independent rates clock model and the LG substitution model (model = 2, aaRatefile = LG.dat; clock = 2). The MCMC analysis was run twice, each with 100,000 generations, discarding the first 50,000 generations as burn-in.

### 2.7 Ancestral character state reconstruction

We selected four morphological traits in the larvae of Trichoptera, including the strategy of respiration, morphology of the anal prolegs, morphology of case or retreat, and habitat. Ancestral character state reconstruction (ACSR) analysis was conducted for each trait, and each feature was individually encoded based on its characteristics (see details in Table S6). Mesquite v3.7.0 (http://mesquiteproject.org) was used to perform maximum likelihood ACSR on deep nodes. The ML reconstruction was conducted under the single-rate “Markov k-state 1 model” (MK1 model). ACSR was performed over 1,000 Bayesian posterior trees of the USCO80_abs75 matrices and summarized on the consensus tree.

## 3. Results

### 3.1 Genome assembly of Trichoptera

The genome size evaluation results indicated that within the suborder Annulipalpia, the genome sizes of Pseudoneureclipsidae and Psychomyiidae were smaller, approximately 179.29 Mb and 177.56 Mb, respectively (Table S7), and the genome sizes of Philopotamidae, Dipseudopsidae, Ecnomidae, Polycentropodidae, and Xiphocentronidae exceeded 200 Mb. In the suborder Integripalpia, notable variation in genome sizes was observed among different families within the “cocoon-maker” group. The family Hydroptilidae exhibited genome sizes ranging from 162.79 to 166.94 Mb. In contrast, Glossosomatidae and Hydrobiosidae displayed larger genomes, approximately 464.4 Mb and 533.62 Mb, respectively. Within the Phryganides, certain species of Phryganeidae, Leptoceridae, Limnephilidae, and Limnocentropodidae exhibited genome sizes exceeding 1 Gb. In addition to the mentioned four families, the genome size assessment for the remaining families within Phryganides indicates sizes larger than 500 Mb and less than 1 Gb. This implied considerable variability in genome sizes within Phryganides as well as larger genomes within these families, possibly associated with their habitat characteristics or other ecological factors influencing their genomic characteristics.

The genomic assessment results, although slightly lower than the actual assembly results, exhibit only a minor difference. The assembly for 86 species of Trichoptera resulted in genome sizes ranging from 124.97 Mb (*Tinodes furcatus* Li & Morse) to 1,353.95 Mb (*Psilotreta porrecta* Yuan, Sun & Yang) in Table S8. The scaffold N50 length ranged from 2.4 kb (*Ecnomus* sp.) to 65.35 kb *Cheumatopsyche brevilineata* (Iwata). The number of scaffolds ranged from 6,821 to 603,762 (*Cheumatopsyche brevilineata* to *Psilotreta porrecta* Yuan, Sun & Yang). The GC content was 26.23%–41.23% (*Dipseudopsis* sp.–*Paduniella communis* Li & Morse). The repetitive sequences accounted for 25%–60% (*Oxyethira* sp.–*Dipseudopsis* sp.;). The Spearman correlation analysis revealed a significant positive correlation between genome size and repetitive sequences in Trichoptera (Spearman correlation coefficient 0.98; *p* < 2.2E16; Fig. S1). Hence, we believed that the augmentation of repetitive sequences was one of the key factors propelling the expansion of trichopteran genomes. The BUSCO completeness assessment, utilizing the insect_odb10 reference dataset, indicated that the genome completeness of the newly sequenced species ranged from 32.00% to 96.50%. Notably, the genome completeness of Annulipalpia was significantly higher than that of Integripalpia (Wilcoxon rank sum test *p* < 0.001). Furthermore, larger genomes and those with higher heterozygosity tend to exhibit lower completeness.

### 3.2 Design of trichopteran USCO set and UCE probe sets

To obtain a high quality USCO dataset for Trichoptera, we used OrthoFinder to assign 157,402 genes from ten trichopteran species and three lepidopteran species to 13,670 orthologous gene families. Of these, 4,181 orthologous gene families were shared by all species, of which 1,682 were identified as single-copy orthologs. We finally obtained a set of 3,860 candidate USCO genes for Trichoptera. These candidate USCO genes were effectively distinguished using a custom-built HMM file, resulting in a final USCO dataset with an average length ranging from 84 to 5,627 bp. Subsequently, we extracted the USCO genes from 111 Trichoptera species using two datasets: the newly developed Trichoptera_odb1 (n=3,860) and Endopterygota_odb10 (n=2,124). By comparing the extraction results of these two datasets, it was evident that the number of single-copy genes acquired from the newly constructed Trichoptera_odb1 was significantly higher than that from the Endopterygota dataset (Fig. S2; *p* > 0.001).

In Annulipalpia, we observed a conspicuous increase in the number of extracted USCO genes for each species within the three superfamilies Hydropsychoidea, Philopotamoidea, and Psychomyioidea. Overall, the species *Cheumatopsyche charites* Malicky & Chantaramongkol (Hydropsychidae) contained the highest number (3,591) of extracted USCO genes (Figs. S3–5). In Integripalpia, a clade within the “cocoon-maker” group, the family Hydroptilidae demonstrated the highest efficiency of USCO gene extraction, with the number of genes ranging from 2,071 to 3,456. However, other families showed a slight improvement in the efficiency of USCO gene extraction, with increases in the number of genes ranging from 289 to 1,143 (Fig. S6). Furthermore, we observed a general increase in the number of extracted USCOs for most species within Phryganides, although a few species in certain families showed a slightly reduced extraction efficiency [(e.g. *Nothopsyche ruficollis* (Ulmer) (Limnephilidae), *Apataniana impexa* Schmid (Apataniidae), *and Triaenodes pelias* Malicky (Leptoceridae); Figs. S7–10]. Finally, based on Trichoptera_odb1 the amino acid sequence lengths of acquired complete USCOs ranged from 84 to 5,638 bp.

For the UCE probe design, we simulated reads ranging from 6,353,682 to 52,208,184 obtained from 17 trichopteran assemblies. These reads were then aligned to the base genome (*Limnephilus lunatus*) in a range of 1.48%–14.89%. Subsequent UCE probe design generated a provisional Trichoptera UCE probe set that included 16,962 bait probes and 9,366 target genes. Finally, 4,792 target genes shared among at least 15 Trichoptera species were selected for design of the final probe set. After removing duplicate probes, we obtained the final Trichoptera UCE probe set (Trichoptera-v1), which consisted of 155,809 baits and 4,731 loci. Based on the trichopteran UCE probe dataset, 4,256 target UCEs were extracted from the genomes of 115 species. The lengths of target UCEs ranged from 242 to 1,739 bp, with most UCE loci falling within the range of 900–1,000 bp (Fig. S11).

### 3.3 Matrix generation

A total of 3,806 USCO genes were obtained from the Trichoptera_odb1. After filtering based on the number of informative sites, we removed 350 genes with fewer than 100 informative sites. 594 genes were removed based on the results of compositional heterogeneity detection, in which the RCV was greater than 0.3, and a total of 2,629 USCO genes were retained following SRH detection. Subsequently, data matrices with taxon occupancy rates ranging of 60%–80% were generated. These matrices included the USCO60/70/80 matrices. IQ-TREE was then used to infer 1,734 gene trees based on the EX_EHO model. For different taxon-occupancy data matrices, genes with an ABS values of each gene tree of >75 were selected, resulting in the creation of the USCO60/70/80_abs75 dataset. Filtered genes were then subjected to additional filtering based on DVMC (<0.8) and treeless (> 0.3) to generate new data matrices, and the relevant details are shown in Table S9.

For UCE loci, a total of 4,256 original UCE markers were obtained using Trichoptera UCE probes in 115 species. UCE markers with fewer than two sequences were removed, resulting in 3,687 UCE markers. Further filtering was performed based on the number of informative sites for each marker, and another 80 additional UCE loci were then removed. The 118 loci with RCV values of 0.15 were excluded based on composition heterogeneity. After the SRH test, 1,366 loci were retained for subsequent matrix generation. Herein, data matrices with taxon occupancy rates of 50%–90% were labeled as UCE50/70/90. Based on the analysis of 1,067 gene trees using GTR model, genes with ABS values of >70 were retained, resulting in the generation of UCE50/70/90_abs70 dataset. These markers were then filtered again based on DVMC (<0.8) and treeless (>0.25), yielding new data matrices. Detailed information regarding each data matrix is provided in Table S9.

### 3.4 Phylogenetic analysis

We used six matrices of the USCO and UCE markers to generate three phylogenetic tree topologies based on different strategies and models. These results showed that all trees supported the monophyly of Trichoptera (Posterior Probabilities ≥95 and SH-aLRT/UFBoot2 ≥ 99). For USCO matrices, topology 1 (T1) was generated using the partitioning and the GHOST model (Figs. 2A; S12–17): (Lepidoptera + (Hydroptilidae + (Annulipalpia + Integripalpia))). In T1, with Hydroptilidae as a sister group to all other Trichoptera, the remaining “cocoon-maker” families formed a monophyletic clade within the suborder Integripalpia, sister to subterorder Phryganides. This topology suggested that the family Hydroptilidae was a basal lineage within Trichoptera, and the traditional distinction between Integripalpia and Annulipalpia was otherwise supported. Simultaneously, we obtained topology 2 (T2) using the MSCM, EX_EHO mixture model, PMSF model, and GAT+GTR model (Figs. 2B, 3; S18–29): (Lepidoptera + (Annulipalpia + Integripalpia)). Here, the family Hydroptilidae was classified in the suborder Integripalpia. It formed a sister group with other Integripalpia, which includes the monophyletic “cocoon-maker” clade and the Phryganides clade. For the UCE matrices, we obtained topology 3 (T3) using UCE matrices based on the MSCM, partitioning model, and the GHOST model (Figs. 2C; S30–38): (Lepidoptera + ((Hydroptilidae + Annulipalpia) + Integripalpia)). In T3, the family Hydroptilidae and Annulipalpia were recovered as sister groups. Overall, the phylogenetic position of the family Hydroptilidae was highly unstable under different makers and models, indicating certain uncertainty in its phylogenetic placement

**Fig. 2.**
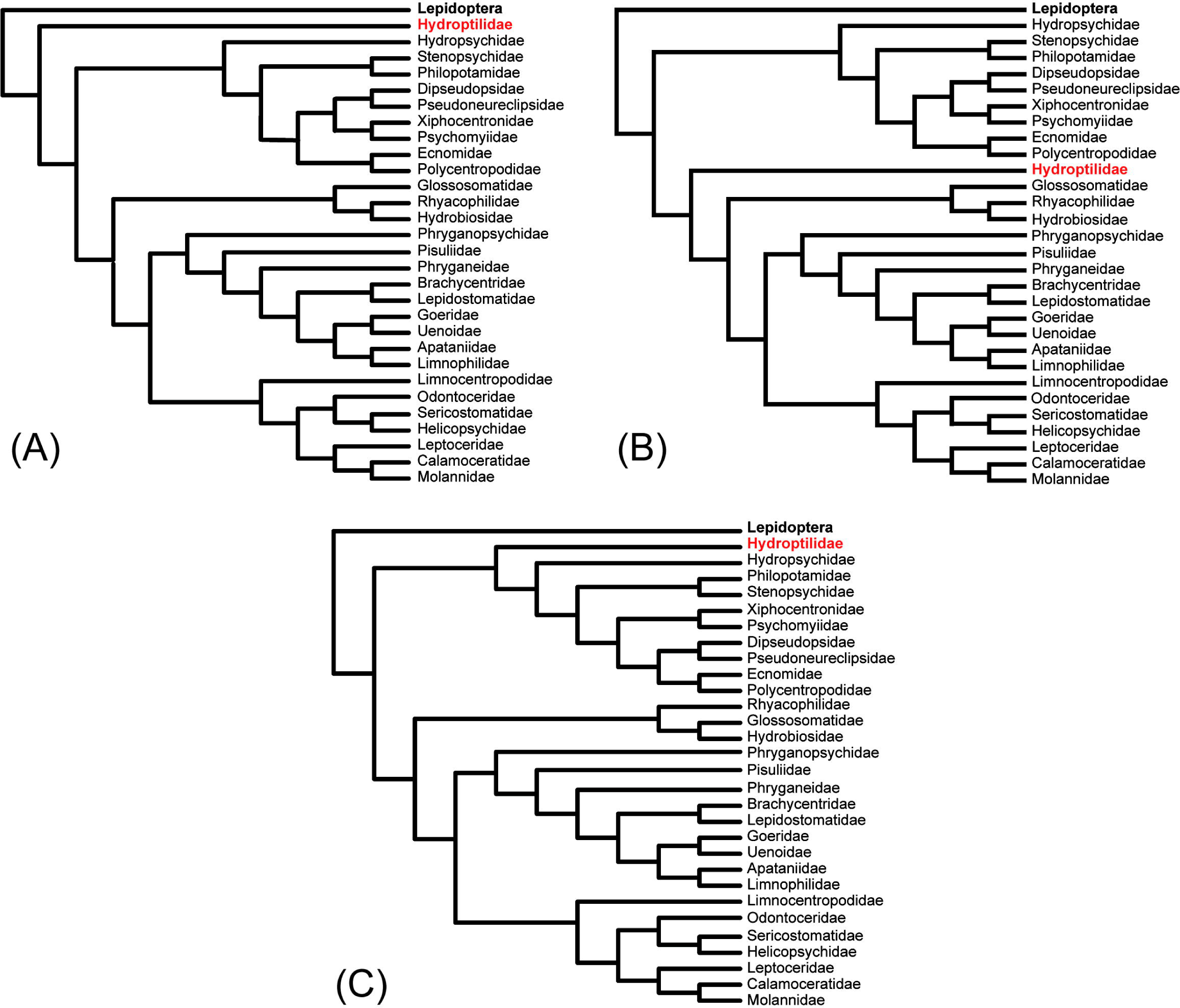
Three topologies of Phylogeny analyses based on USCO and UCE. A: USCO70/80_abs75 matrices based on the partitioning model and the GHOST model, USCO60 matrices based on the GHOST model; B: USCO60 matrices based on partitioning model, and USCO60/70/80_abs75 matrices based on the EX_EHO mix model, PMSF model, MSCM and Dayoff6 recoding; C: UCE50/70/90_abs70 matrices based on the partitioning model and the GHOST model.

**Fig. 3.**
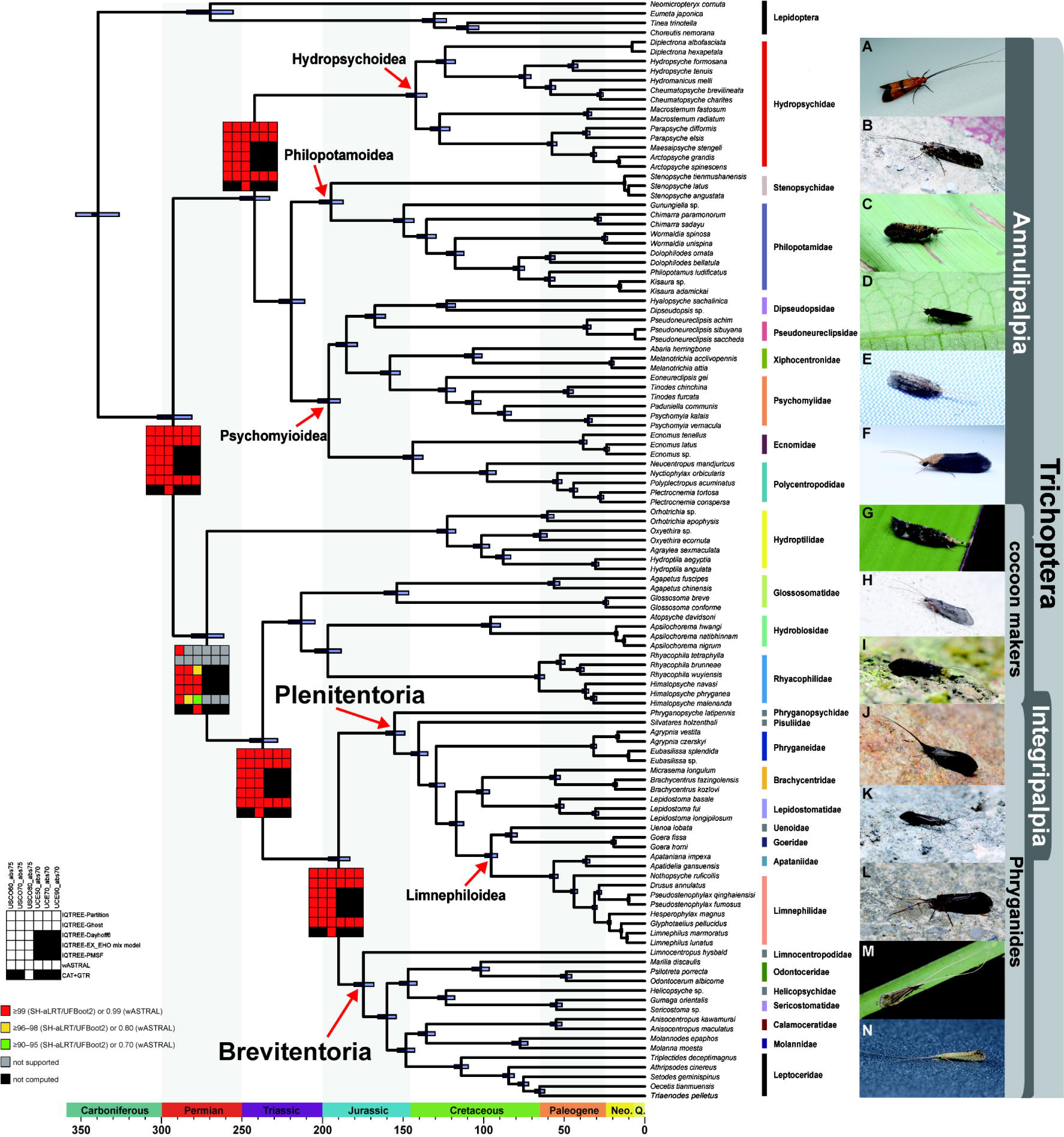
Phylogeny and divergence time of Trichoptera inferred from matrix USCO80_abs75 using the CAT+GTR model implemented in PhyloBayes. Node supports from all analyses are indicated by the colored squares. The gray squares are shown for the nodes inconsistent with BI tree. Node bars represent 95% confidence intervals (CIs) of the estimated divergence times integrated from all MCMC runs. A: *Macrostemum fastosum* (Hydropsychidae); B: *Stenopsyche grahami* (Stenopsychidae); C: Philopotaminae sp. (Philopotamidae); D: *Melanotrichia* sp. (Xiphocentronidae); E: *Ecnomus tenellus* (Ecnomidae); F: *Nyctiophylax* sp. (Polycentropodidae); G: (k) *Hydroptila* sp. (Hydroptilidae); H: Glossosomatidae sp. (Glossosomatidae); I: *Rhyacophila* sp. (Rhyacophilidae); J: *Lepidostoma* sp. (Lepidostomatidae); K: *Apatidelia egibie* (Apataniidae); L: *Pseudostenophylax kriton* (Limnephilidae); M: Odontoceridae sp. (Odontoceridae). N: *Setodes* sp. (Leptoceridae).

In the suborder Annulipalpia, the monophyly of Hydropsychoidea, Philopotamoidea, and Psychomyioidea was strongly supported, with a classical phylogenetic relationship (Hydropsychoidea + (Philopotamoidea + Psychomyioidea)) observed for all three topologies (Fig. 2). Within Psychomyioidea, both loci produced topologies that recovered the sister group relationships between Dipseudopsidae and Pseudoneureclipsidae, Psychomyiidae and Xiphocentronidae, and Ecnomidae and Polycentropodidae. However, these three clades showed different relationships in the three topologies.

In Integripalpia, the other “cocoon-maker” families (i.e., excluding Hydroptilidae) formed a monophyletic group in all topologies. According to the USCO phylogenetic tree, this monophyletic group is recovered as (Glossosomatidae + (Hydrobiosidae + Rhyacophilidae)), although the UCE tree did not exhibit the same result, instead yielding ((Glossosomatidae +Hydrobiosidae) + Rhyacophilidae). However, the UCE result did not have good support. In Phryganides, the relationships within the infraorder Plenitentoria were strongly supported, and the monophyly of Limnephiloidea was recovered. Within the infraorder Brevitentoria, the family Odontoceridae was ambiguously classified as Sericostomatoidea (Fig. 3). Further topology testing indicated that T2 was strongly supported and T1 was strongly rejected in all tests (AU, WKH, and WSH; logL: - 9365528.003; *p* < 0.05; Table S10). This indicated that the family Hydroptilidae should be treated as a basal clade of Integripalpia rather than an independent basal clade within Trichoptera. Similarly, T3 was rejected in all hypothesis tests, suggesting that Hydroptilidae is not a member of the suborder Annulipalpia.

### 3.6 Divergence time and ancestral state reconstruction

Divergence time was estimated using the BI tree based on the USCO80_abs75 dataset with calibration performed based on 12 fossil calibration points. These results indicated that the origins of Trichoptera occurred during the Early Permian, around 281.16–302.52 Ma (95% Highest Posterior Density, HPD; Fig. 3). The time of origin of the suborder Integripalpia preceded that of the suborder Annulipalpia, and the divergence of these two suborders occurred within these two distinct periods. The origin of Integripalpia occurred during the Middle Permian period (261.39– 281.32 Ma), whereas Annulipalpia originated during the Early Triassic (231.94–251.1 Ma). In the suborder Annulipalpia, the origins of Hydropsychoidea, Philopotamoidea, and Psychomyioidea all occurred during the Early Jurassic–Cretaceous period (i.e., 135.31–148.08 Ma, 187.25–201.94 Ma, 189.09–202.9 Ma, respectively). Divergences between the ancestor of the other “cocoon-maker” groups (Glossosomatidae, Hydrobiosidae, Rhyacophilidae) and Phryganides, and between Plenitentoria and Brevitentoria occurred during the Triassic–early Jurassic (i.e., 227.95–245.1 Ma and 183.07–196.35 Ma, respectively).

ACSR showed that the common ancestor of Trichoptera exhibited through undifferentiated integument (Figs. S39–40). A combination of the gill and integument respiration was inferred to be independent synapomorphies of Phryganides, Hydropsychidae, and *Himalopsyche* (Fig. 39a). Moreover, the species living in flowing water were commonly accompanied by strongly developed anal prolegs (Figs. S39b, 40b). The free-living state, without any type of shelter (e.g., similar to early instars of Hydroptilidae), was probably the ancestral state for Trichoptera and, within Integripalpia, the hydroptilid purse case, the glossosomatid saddle case, and Phryganides tube case appear to have evolved independently from the ancestral pattern (Fig. S40a).

## 4. Discussion

### 4.1 Genomic data and maker development

This study assembled genomes covering 28 families within Trichoptera. The sizes of assembled genomes ranged from 124.97 Mb to 1,353.95 Mb, with the largest assembly being approximately 11 times larger than the smallest. Specifically, the genomes of the families Psychomyiidae, Xiphocentronidae, and Pseudoneureclipsidae were smaller than the known minimum genome size in Trichoptera, represented by *Agraylea sexmaculata* Curtis (196.07 Mb; Heckenhauer et al., 2022). Here, most species within the suborder Integripalpia generally exhibited larger size genomes sizes than the majority of taxa in the suborder Annulipalpia, thus further confirming previous findings (Heckenhauer et al., 2022). Phylogenomics has been successfully applied to phylogenetic studies of various arthropod taxa on levels ranging from order to species (Bossert et al., 2019; Bradford et al., 2022; Buenaventura et al., 2021; Johnson et al., 2022; Stephen. et al., 2017; Yu et al., 2022; Zhang et al., 2022). Moreover, it has been employed in population genomics (Winker et al., 2018). Within Hexapoda, taxa with available USCO datasets include Collembola (n = 1,997), Lepidoptera (n = 5,286), Hymenoptera (n = 5,991), Hemiptera (n = 2,510), Diptera (n = 3,285), and Endopterygota (n = 2,124) (Waterhouse et al., 2018). In addition, UCE probe sets are available for Coleoptera, Diptera, Lepidoptera, Hemiptera, and Hymenoptera (Branstetter et al., 2017; Faircloth, 2017).

The number of USCO loci included in the novel designed dataset was notably higher than that in the USCO dataset for Holometabola, as reported by Waterhouse et al (2018). This increase in loci number may contribute to a more comprehensive understanding of the phylogenetic relationships within Trichoptera. Our results also revealed that when extracting USCO genetic data from Trichoptera datasets, especially for species with smaller genomes within the suborder Annulipalpia, the quality of the resulting assembly significantly improved. This improvement was particularly notable for species with high quality genomes assembled using PacBio or Oxford Nanopore long-read sequencing technologies. Moreover, although our results have significant limitations in terms of number and average length compared with the data obtained from Lepidopteran USCO datasets, continuous improvement and addition of new high-quality genome assemblies may enhance the quality of the USCO dataset over time. The trichopteran UCE probe set designed in this study had a higher number of bait probes and UCE loci (Table S11). In contrast to USCO, UCE probes exhibited less variation in extraction efficiency between the two suborders and showed more consistent performance across groups with larger genomes within Integripalpia. This phenomenon has also been observed in studies involving other arthropod groups (Zhang et al., 2019). Considering the larger genomes within Integripalpia, UCE probes can be used to extract more phylogenetic information sites, thereby mitigating the impact of assembly fragmentation when conducting phylogenetic studies. With the increasing prevalence of full-length sequencing technologies and rising number of high-quality trichopteran genome assemblies and transcriptomes, newly designed USCO and UCE datasets can be used to extract more effective information for further phylogenetic analyses of Trichoptera.

### 4.2 Which is best supported topology?

During the reconstruction of phylogenetic relationships, issues such as compositional heterogeneity, inclusion of paralogs, and assembly errors can introduce systematic errors. In addition, errors in multiple sequence alignment, and excessive trimming can lead to weakening of phylogenetic signals (Ashkenazy et al., 2018; Steenwyk et al., 2020). To avoid treatment errors, we used different strategies to screen the loci extracted by two markers. Subsequently, we also used multiple models to reconstruct the higher taxonomic relationships within Trichoptera. The phylogenetic relationship of most groups was strongly supported under different models. However, the phylogenetic positions of a few families showed conflicting results among the topologies generated by different molecular markers, with the Hydroptilidae position being particularly unstable.

The analysis of gCF and sCF indicated that the inconsistency in gene trees was the primary contributor to the systematic errors observed in our phylogenetic reconstruction (Figs; S12–14, S18, S22, S30–32; Salichos and Rokas, 2013). USCO70/80 matrices based on the partitioning and the GHOST model, and USCO60 matrice based on the GHOST model generated T1. This topology contradicts all previous studies based on morphology and a few markers in phylogeny. Furthermore, T1 was rejected by the topology test (Table. S9). We believe that the issues with this topology may be due to systematic error. A comparison of traditional substitution models, MSCM and site-heterogeneous models (i.e., PMSF and CAT+GTR) can effectively mitigate systematic errors caused by ILS and LBA phylogenetic reconstruction and has been found to help resolve the phylogenetic positions of several anciently diverged lineages (Galindo et al., 2021; Wang et al., 2017; Zhang et al., 2018). Herein, the results obtained from phylogenetic inference using MSCM, PMSF, and CAT+GTR strongly supported the classification of Hydroptilidae as a basal clade within the suborder Integripalpia (T2). This hypothesis is consistent with the results of previous studies based on morphological evidence or marker-based phylogenies (Kjer et al., 2016; Ross, 1956,1967; Thomas et al., 2020); moreover, the topology tests also showed a strong preference for T2. Simultaneously, compared with the concatenation-based, site-homogeneous, and the GHOST model, the EX_EHO mixture model was effective in reducing systematic errors, resulting in more plausible phylogenetic inferences (Fig. S18–20; Feuda et al., 2017; Marlétaz et al., 2019; Williams et al., 2020). In general, substitutional saturation by ancient rapid divergence can lead to incongruence and inaccurate phylogenetic inferences (Laumer et al., 2018). Our results suggest that these issues can be addressed using amino acid recoding. Notably, the partial node of the ML tree determined based on USCO80 using Dayhoff6 recording was not well-supported (Fig. S25), indicating that amino acid recoding can also lead to the erosion of phylogenetic information (Foster et al., 2022).

We observed significant differences between the trees generated by UCE and USCO marker sets. All phylogenetic trees based on UCE constructed using the partitioning, GHOST, and MSCM models showed that Hydroptilidae is a member of the suborder Annulipalpia, which is consistent with the hypothesis proposed by Ge (2023). Notably, the phylogenetic relationships generated within the infraorder Brevitentoria based on UCE markers using MSCM remain unclear (Figs. S35– 37). These results contradict those of previous studies based on morphology and multiple molecular markers (Johanson et al., 2017). Furthermore, topology testing also suggested that T3 may be inaccurate. UCE markers may be influenced by issues such as compositional biases or model violation, both of which can lead to inaccurate phylogenetic reconstruction (Baker et al., 2021). Therefore, we suggest that the applicability and reliability of UCE markers in the analysis of trichopteran lineages warrant further investigation.

### 4.3 Phylogeny of Trichoptera

Since Martynov’s system was proposed a hundred years ago, the phylogenetic position of the “cocoon-maker” group has been controversial (Ge et al., 2023; Ivanov, 2002; Kjer et al., 2016; Ross, 1967; Schmid, 1998; Wiggins and Wichard, 1989). The primitive morphological characteristics (i.e., campodeiform larvae and semipermeable cocoon) and living behaviors (i.e., free-living) of “cocoon-maker” larvae have influenced researcher speculation regarding their phylogenetic placement.

Ptilocolepidae is a small family closely related to Hydroptilidae, and was formerly considered a subfamily within this clade (Malicky, 2001). Since its elevation to family status, the phylogenetic position of Ptilocolepidae has been contentious (Malicky, 2008; Thomas et al., 2020; Thomson et al., 2022). In the reconstructed phylogenetic relationships of Trichoptera based on USCO, the topology with the best placement of Hydroptilidae indicated that it is sister to all other Integripalpia. Although we did not collect specimens of Ptilocolepidae, based on previous molecular and morphological studies, we suggest that Ptilocolepidae and Hydroptilidae should be classified as Hydroptiloidea and be considered as the basal lineages within Integripalpia. This result would be consistent with hypotheses based on analyses of 18S/28S ribosomal RNA genes and other molecular markers (Kjer et al., 2016; Thomas et al., 2020). Compared to previous studies of the phylogenetic relationships of Trichoptera based on relatively few nuclear and mitochondrial markers, the phylogenetic positions of Glossosomatidae, Hydrobiosidae, and Rhyacophilidae were stably recovered by the analyses reported here. We found that they form a monophyletic clade as a sister group to the Phryganides in phylogenetic reconstruction using both USCO and UCE markers. This phylogenetic relationship also appears in the study of Frandsen et al (2023). We therefore suggest that these three families should be classified as Rhyacophiloidea (Fig. 3). We also note that their last larval instar builds a fixed, dome-like pupal case of stones and silk, which also suggests a common morphological origin (Morse et al., 2019; Wiggins, 2004; Wiggins and Wichard, 1989).

The phylogenetic relationship within Psychomyioidea based either on morphology, a few markers, and the mitogenome is ambiguous, especially for Pseudoneureclipsidae (Chamorro and Holzenthal, 2011; Johanson and Espeland, 2010; Johanson et al., 2012; Thomas et al., 2020). For example, based on synapomorphies, the morphological characteristics of female sternum VIII sclerites, and the presence of a long larval spinneret without labial palpi, Pseudoneureclipsinae should be placed in Dipseudopsidae a sister clade to Dipseudopsinae (Li et al., 2001). Combined with Tachet’s (2010) study on the shelter shape of Pseudoneureclipsidae, our phylogenetic results substantiate the hypothesis proposed by Li et al. (2001), and strongly support the sister group relationship between Pseudoneureclipsidae and Dipseudopsidae, which are more closely related to the sister branches formed by Psychomyiidae and Xiphocentronidae. We also consider that previous mitochondrial studies may reflect the fact that similar rearrangements of mitochondrial structure may be the result of convergent evolution (Greenway et al., 2020). The phylogenetic relationships reported here also suggest that apomorphy protein-coding gene rearrangements of Pseudoneureclipsidae, Ecnomidae, and Polycentropodidae may not serve as effective phylogenetic markers.

Within Integripalpia, our phylogenetic analysis recovers paraphyletic Phryganeoidea and monophyletic Limnephiloidea lineages. However, the monophyly of Sericostomatoidea and Leptoceroidea is not supported. Specifically, Odontoceridae is classified as Sericostomatoidea, a relationship that also aligns with the findings of Malm et al. (2013). However, due to the scarcity of specimens and genomic information from other families within Sericostomatoidea, determination of the precise phylogenetic position of Odontoceridae and the questions whether Odontoceridae belongs to Leptoceroidea warrant further investigation involving a broader array of taxonomic units.

### 4.4 Origin and adaptive evolution of Trichoptera

Divergence time analyses revealed that the most recent common ancestor of Trichoptera occurred in the Early Permian period (approximately 292 Ma), and the divergence time of Trichoptera is earlier than that reported by previous studies (Malm et al., 2013; Thomas et al., 2020, 2023). ACSR suggests that the ancestors of all extant Trichoptera most likely lived in flowing water, similar to the Annulipalpia clade. We speculate that subsequent differentiation was strongly correlated with habitat change. In other words, adaptations to life in lentic waters and in slowly moving water likely evolved independently across integripalpian and annulipalpian lineages.

The ancestors of the suborder Annulipalpia evolved into running water, where fast flowing water resulted in less sediment and provided a constant supply of dissolved oxygen and food. This led to the retention of well-developed anal prolegs and construction of fixed shelters, thereby reducing larval movement disturbance and providing better camouflage defenses against predators (Morse et al., 2019; Wiggins, 2004). The adaptation of the Psychomyioidiea to slowly moving water has promoted the evolution of filtering nets into capture nets to obtain food more efficiently. In the Integripalpia, three cases (purse-case, saddle-case, and tubular cases) provide better protection for the larvae, while cocoon making preserves the relatively primitive purse and saddle cases. The larvae of microcaddisflies (Hydroptilidae and Ptilocolepidae) have rapidly developing first four free-living instars and the final (fifth) instar larva constructs various types of purse-case. Dissimilarly, each instar of glossossomatid larvae constructs a saddle-case, similar to the pupal shelters of Hydrobiosidae and Rhyacophilidae, unlike the tube cases constructed by Phryganides (Wiggins, 2004). The phylogenetic relationships of these taxa provide evidence of evolution from purse-cases to saddle-cases to tubular cases and the saddle-case and tubular cases have a common origin. The phylogenetic relationships of these taxa provide evidence of evolution from purse-cases to saddle-cases to tubular cases and the saddle-case and tubular cases have a common origin. In Rhyacophiloidea, sister families Rhyacophilidae and Hydrobiosidae in which the last larval instar also builds a fixed, dome-like pupal case. However, it’s noteworthy that their larvae remain free-living in all instars. Consequently, evolutionarily speaking, they discarded the larval case for easier predatory mobility (Thomas et al., 2020).

The common ancestor of Phryganides originated in the Early Jurassic. Frakes (1979) and Hallam (1994) suggested that the breakup of the Pangaea supercontinent, the formation of new ocean basins, and changes in ocean circulation patterns during the Late Triassic led to climate changes in many areas. During the mid-to-late Jurassic period, the frequency of emergent xerophytes in northern Chile and southern parts of Russia increased in desert areas, further supporting this inference (Hartley et al., 2005; Vakhrameev, 1964). Some studies have shown that climate change can also affect water flux (Markovic et al., 2017). During this period, the decrease in flow flux in some areas led to higher levels of bottom sediment, which may have increased the hydrological diversity. In general, the static water environment increased pressure on the caddisflies to obtain oxygen and food and promoted they to use more materials (i.e., grit, leaf fragments, and decomposing bark) to build portable cases to protect themselves, find food, and create water flow to obtain dissolved oxygen (Wiggins, 1996). Thus, the breakup of Pangaea may be one of the factors promoting the diversification and evolution of Phryganides. Exploring the relationship between species divergence and paleoenvironmental events in Trichoptera and other aquatic insects may be an interesting topic for future phylogenomic studies. This could be achieved by improving taxon sampling and incorporating new fossil evidence. We can gain a deeper understanding of the evolutionary processes and adaptations of these insect groups in response to environmental changes over time.

## CRediT authorship contribution statement

X.G., C.S. and B.W. conceived and designed the experiments. X.G. L.P. and H.Z collected the samples. X.G. analyzed the data and results. X.G. wrote the manuscript. X.G. and J.W. produced diagram. X.G., J. C.M., L.Y., C.S. and B.W. revised the manuscript. All authors read and approved the final manuscript.

## Acknowledgements

We sincerely thank the editors and reviewers for their valuable comments on this study. We greatly thank Prof. Feng Zhang (Nanjing Agricultural University) and Prof. Liang Lv (Hebei Normal University) for their valuable suggestions on the phylogenetic analyses. We also thank Dr. Zhen-xing Ma, De-wen Gong (both Nanjing Normal University) and Dr. Qing-bo Huo (Yangzhou University) for collecting some samples. Thanks also are extended to Mr. Qian-le Lu for providing photos of some sequenced samples. This research was supported by the National Natural Science Foundation of China (32271631; 32311520285).

## Data accessibility

The matrix, USCO dataset, UCE probe, ML tress, and BI trees are available on Zenodo DOI: 10.5281/zenodo.10634334. The newly assembly genomes are available at NGDC (BioProject ID: PRJCA022723), and the accession numbers are available Table S2.

