## Supplementary data figure for "Phylogenomics resolves a 100-year-old debate regarding the evolutionary history of caddisflies (Insecta: Trichoptera)"

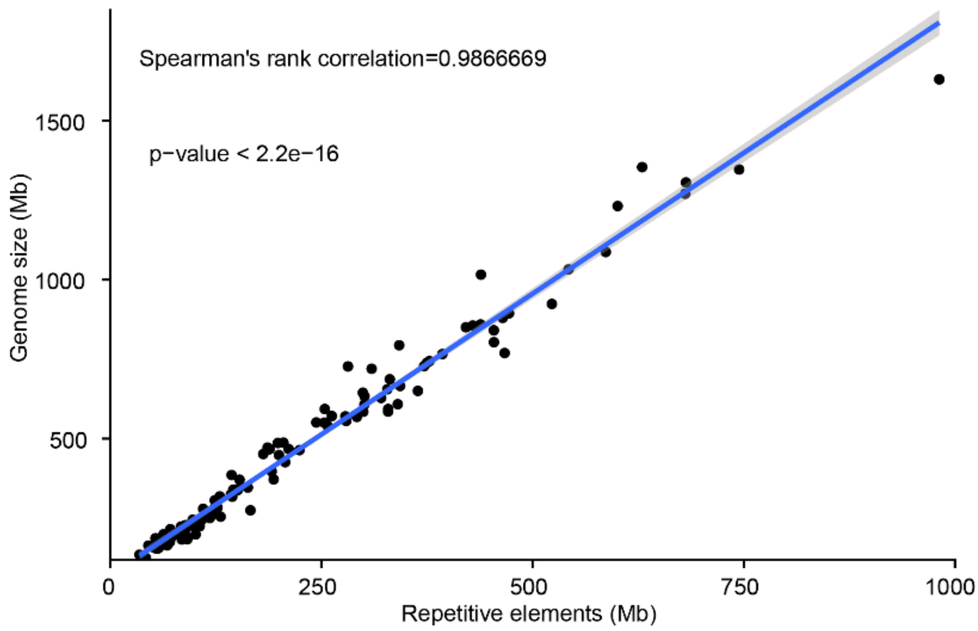


**Fig. S1** Spearman's rank correlation between genome size and repetitive elements.

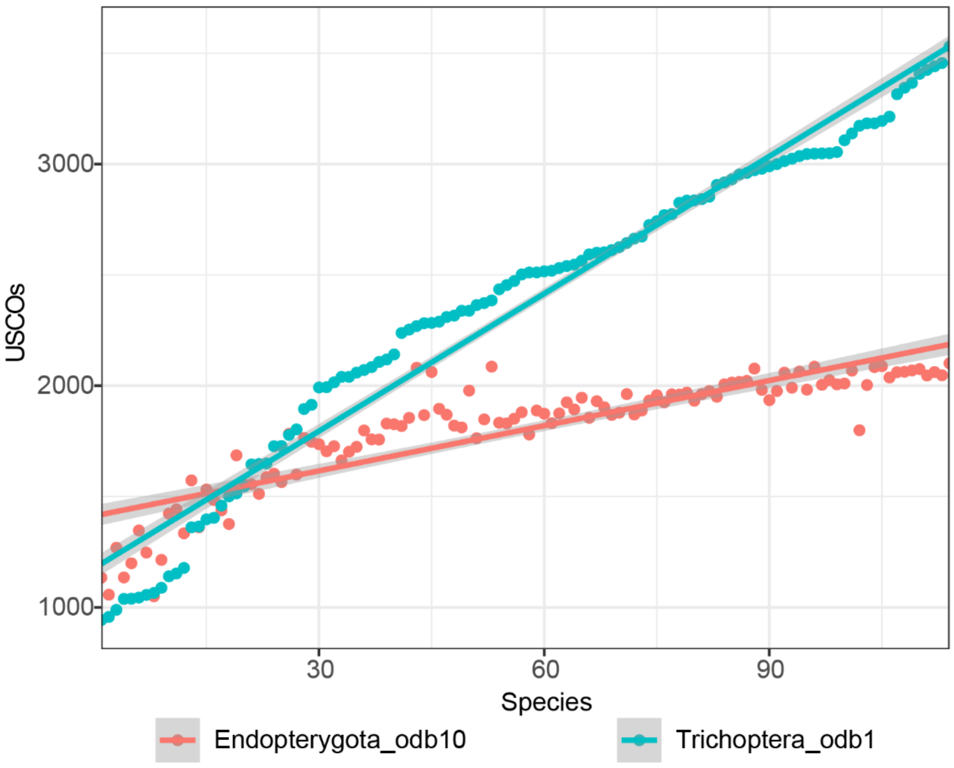


**Fig. S2** Comparison of the USCO extracted from the Trichoptera genome based on two data. datasets

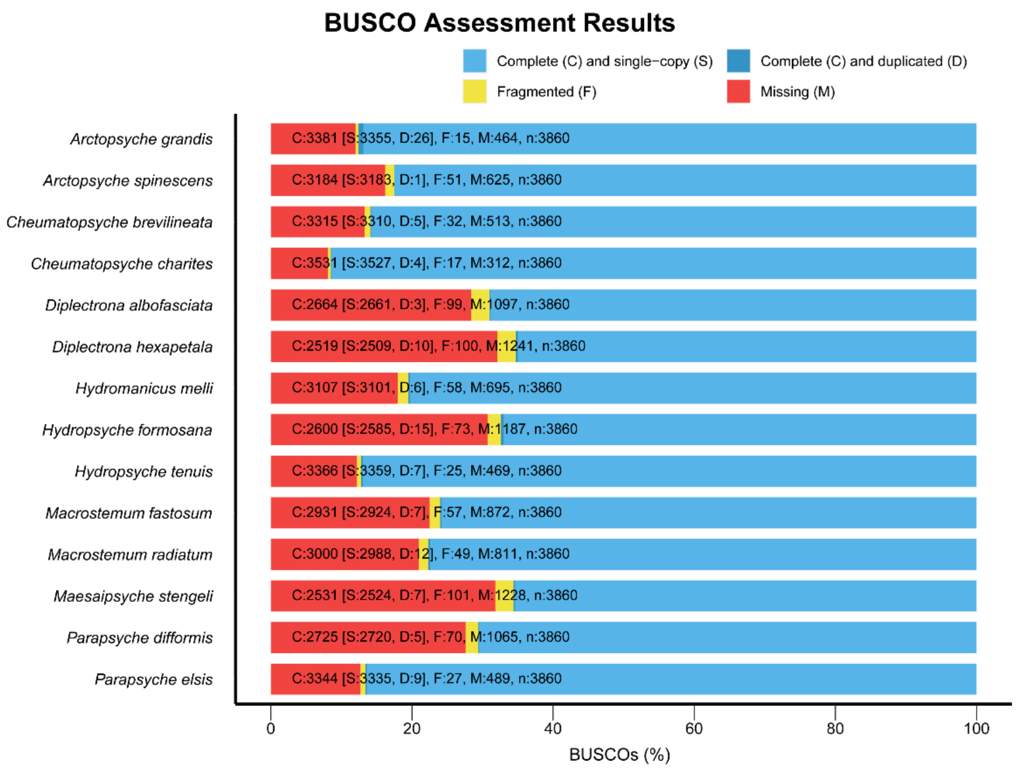


**Fig. S3** BUSCO genome completeness assessments of Hydropsychoidea: complete (C, blue), complete single-copy (S, light blue), complete duplicated (D, dark blue), fragmented (F, yellow), and missing (M, red).

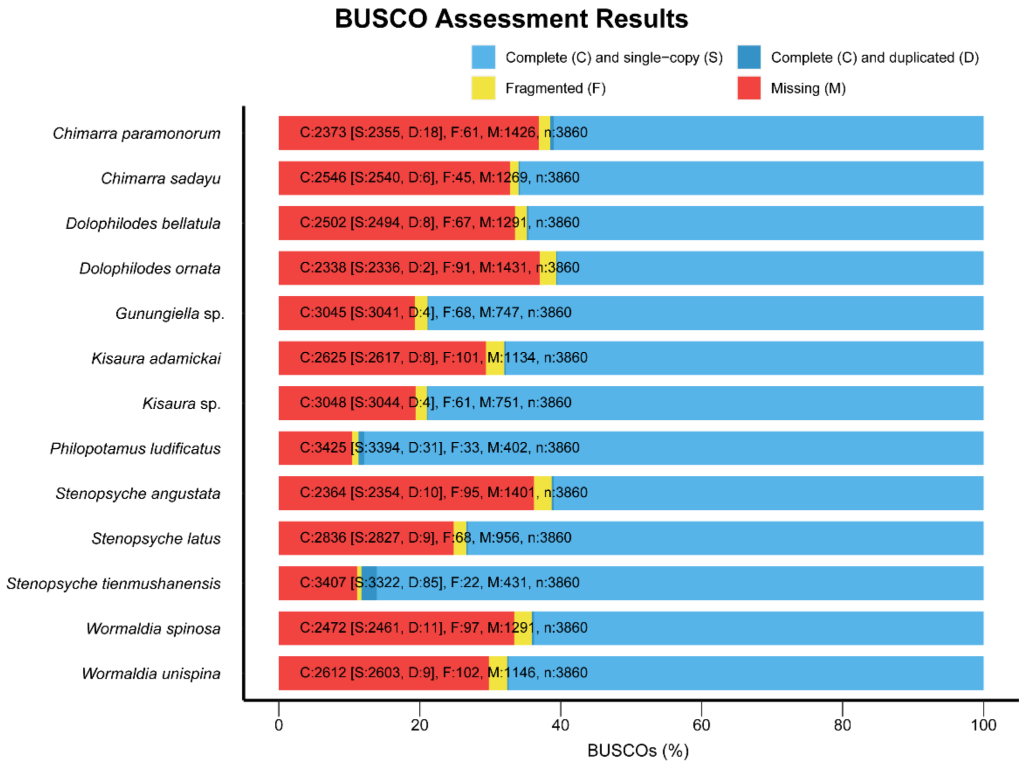


**Fig. S4** BUSCO genome completeness assessments of Philopotamoidea: complete (C, blue), complete single-copy (S, light blue), complete duplicated (D, dark blue), fragmented (F, yellow), and missing (M, red).

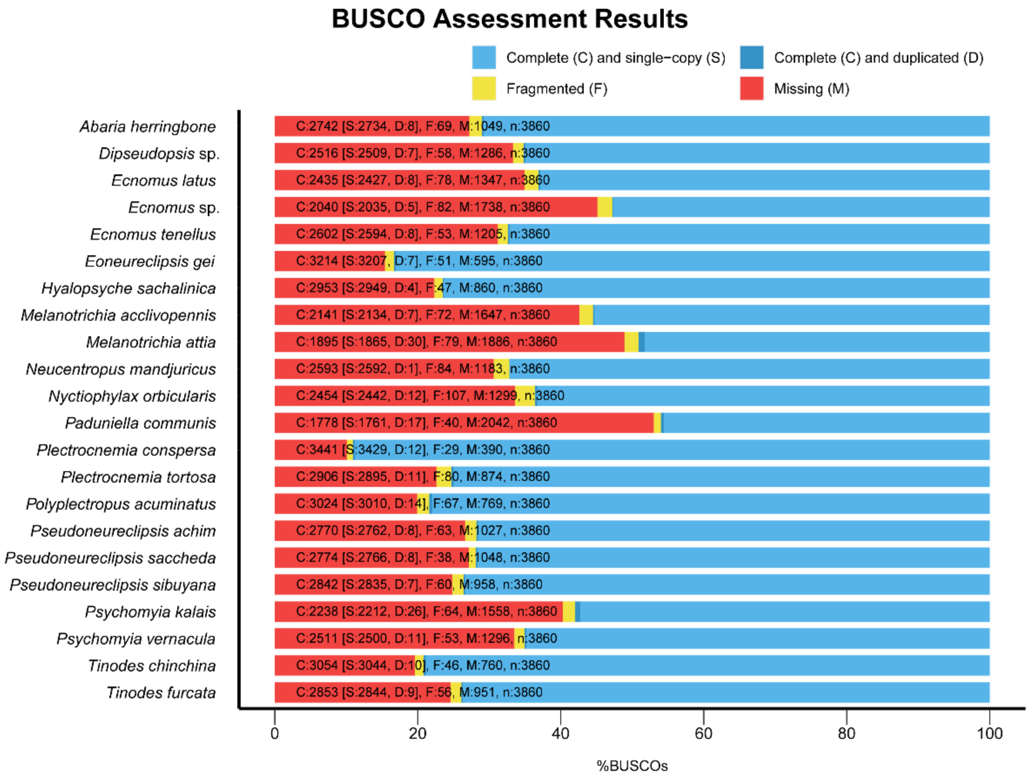


**Fig. S5** BUSCO genome completeness assessments of Psychomyioidea: complete (C, blue), complete single-copy (S, light blue), complete duplicated (D, dark blue), fragmented (F, yellow), and missing (M, red).

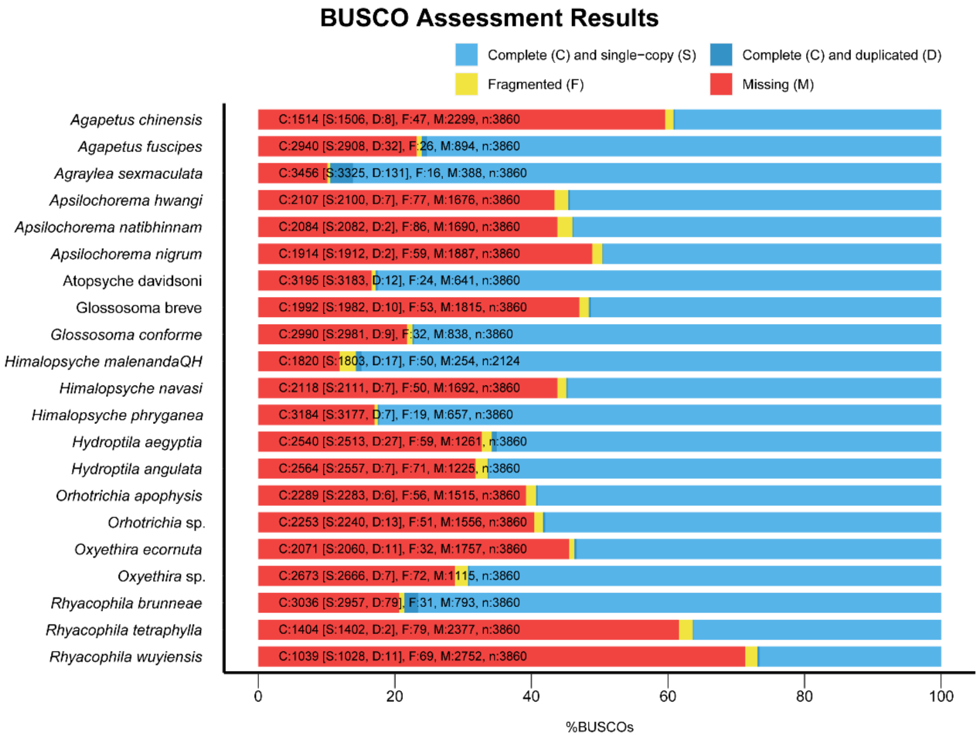


**Fig. S6** BUSCO genome completeness assessments of cocoon-maker: complete (C, blue), complete single-copy (S, light blue), complete duplicated (D, dark blue), fragmented (F, yellow), and missing (M, red).

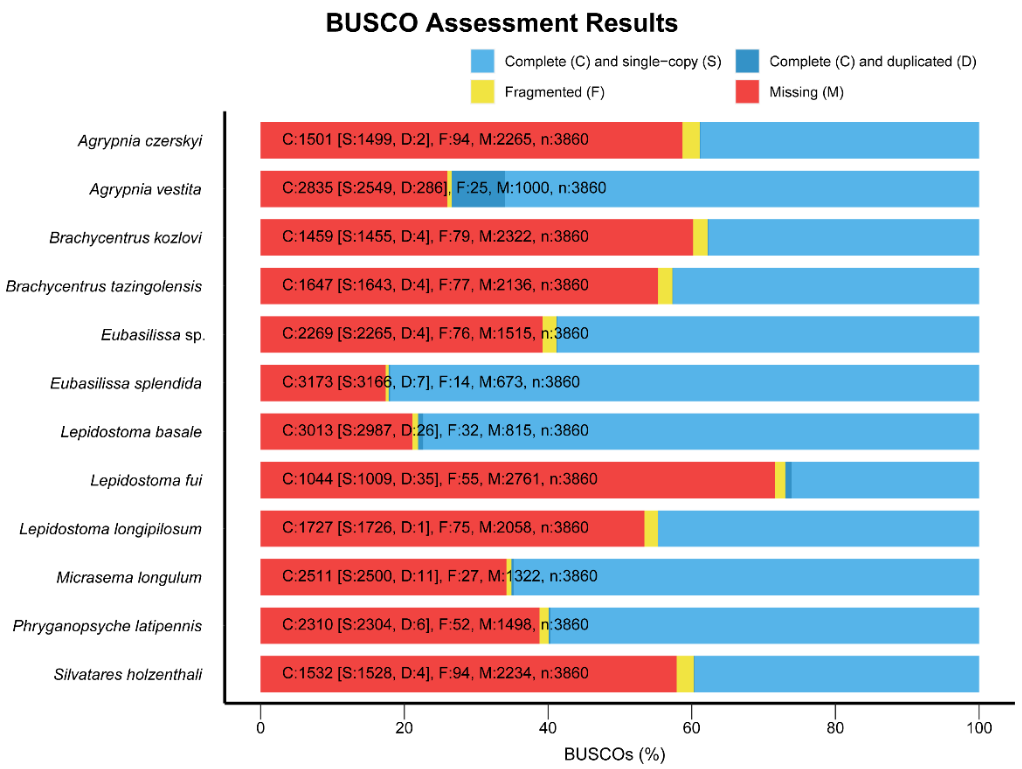


**Fig. S7** BUSCO genome completeness assessments of Phryganeoidea: complete (C, blue), complete single-copy (S, light blue), complete duplicated (D, dark blue), fragmented (F, yellow), and missing (M, red).

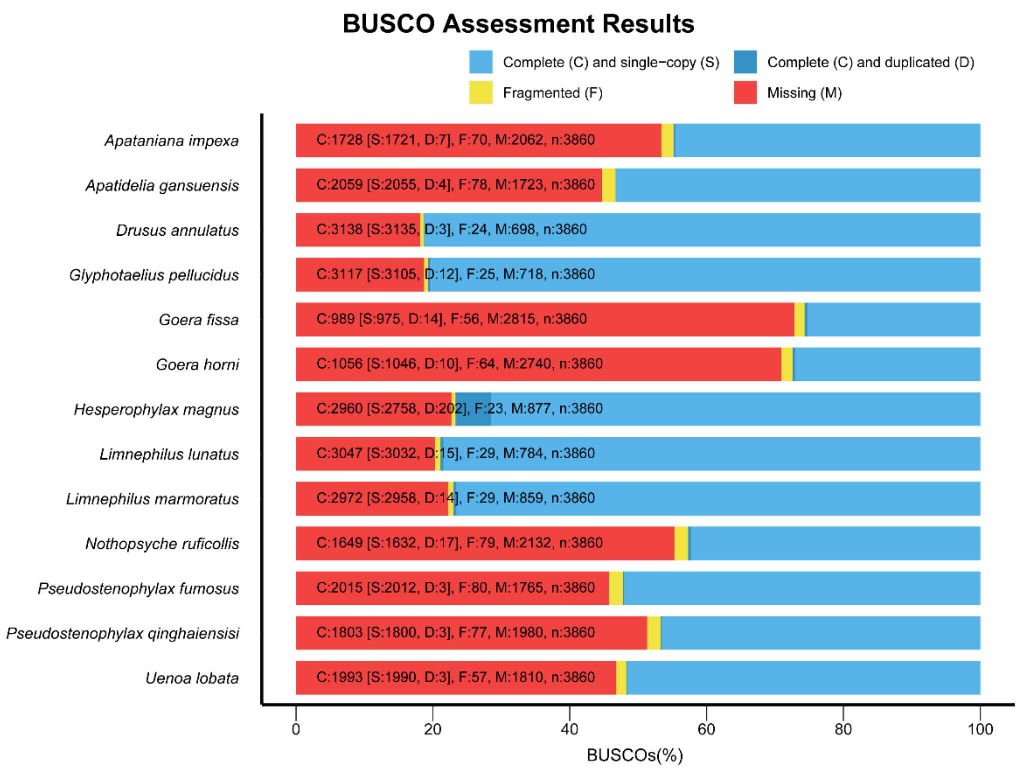


**Fig. S8** BUSCO genome completeness assessments of Limnephiloidea: complete (C, blue), complete single-copy (S, light blue), complete duplicated (D, dark blue), fragmented (F, yellow), and missing (M, red).

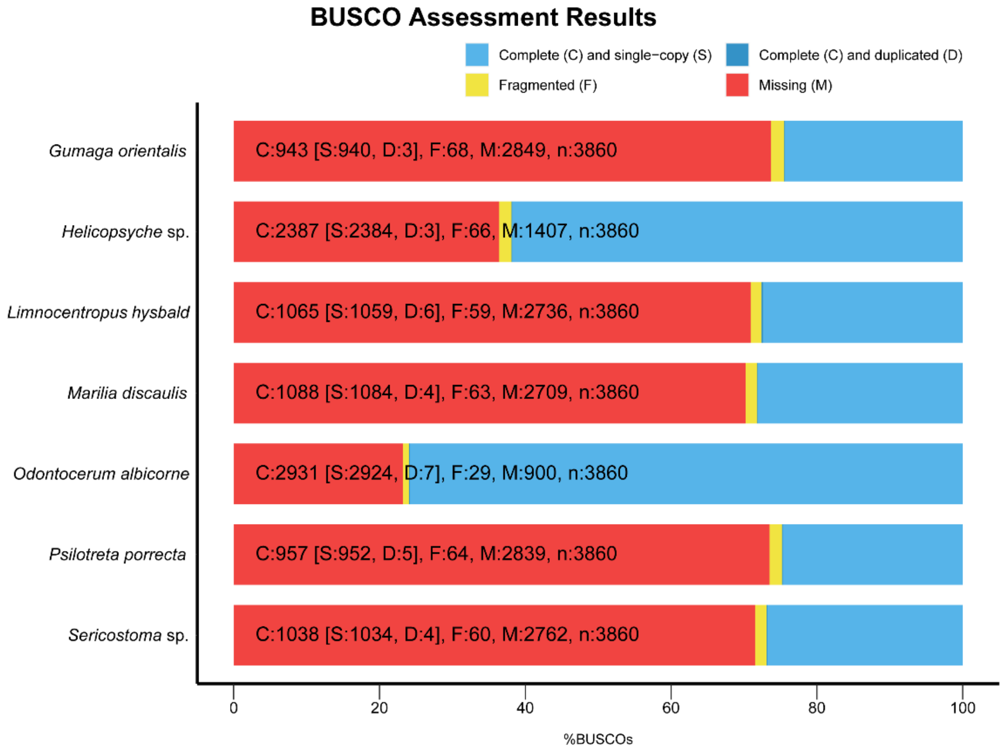


**Fig. S9** BUSCO genome completeness assessments of Sericostomatoidea: complete (C, blue), complete single-copy (S, light blue), complete duplicated (D, dark blue), fragmented (F, yellow), and missing (M, red).

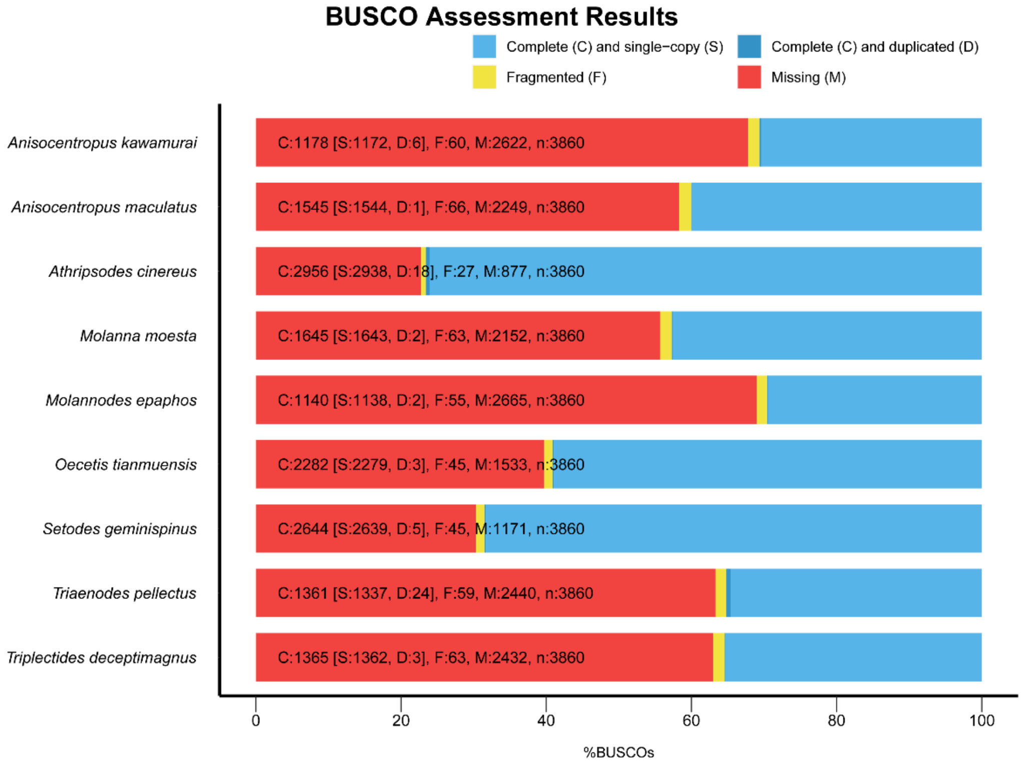


**Fig. S10** BUSCO genome completeness assessments of Leptoceroidea: complete (C, blue), complete single-copy (S, light blue), complete duplicated (D, dark blue), fragmented (F, yellow), and missing (M, red).

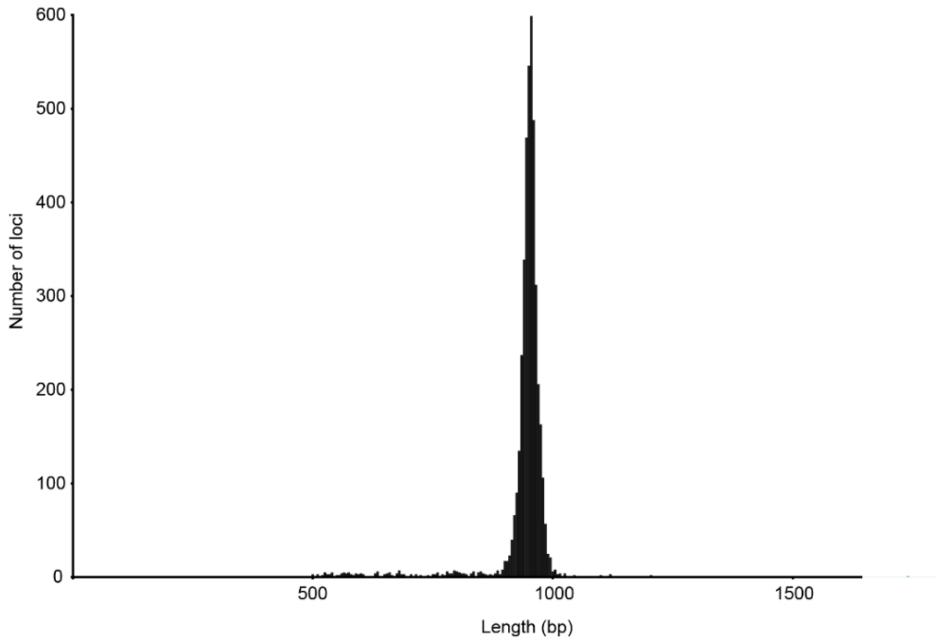


**Fig. S11** Extraction marker length of 111 Trichoptera based on UCE probe set of Trichoptera.

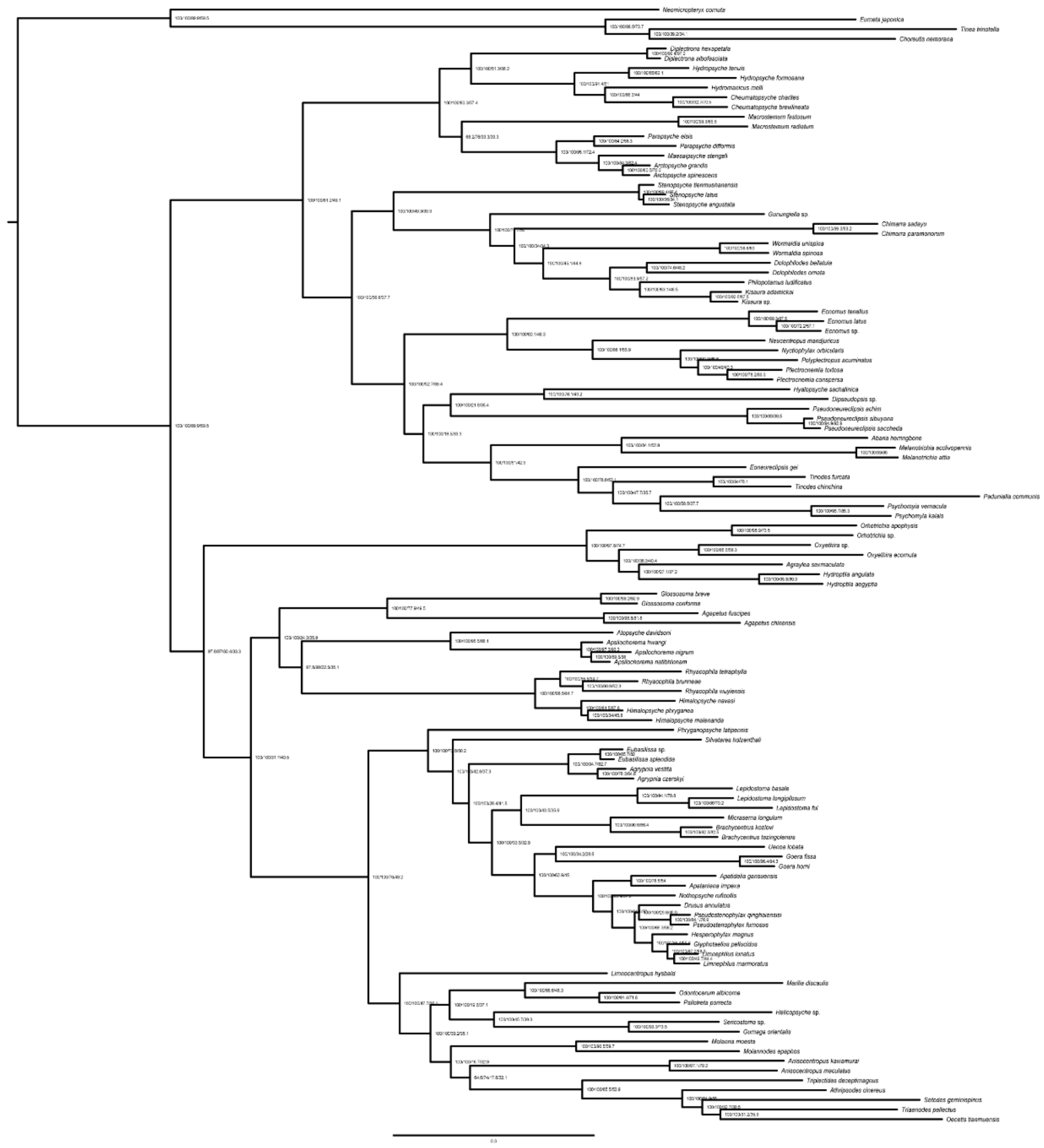


**Fig. S12** ML phylogenomic tree of Trichoptera based on USCO60_abs75 dataset with Partitioning model in IQ-TREE. Node values represent SH-aLRT/UFBoot2/gCF/sCF, respectively.

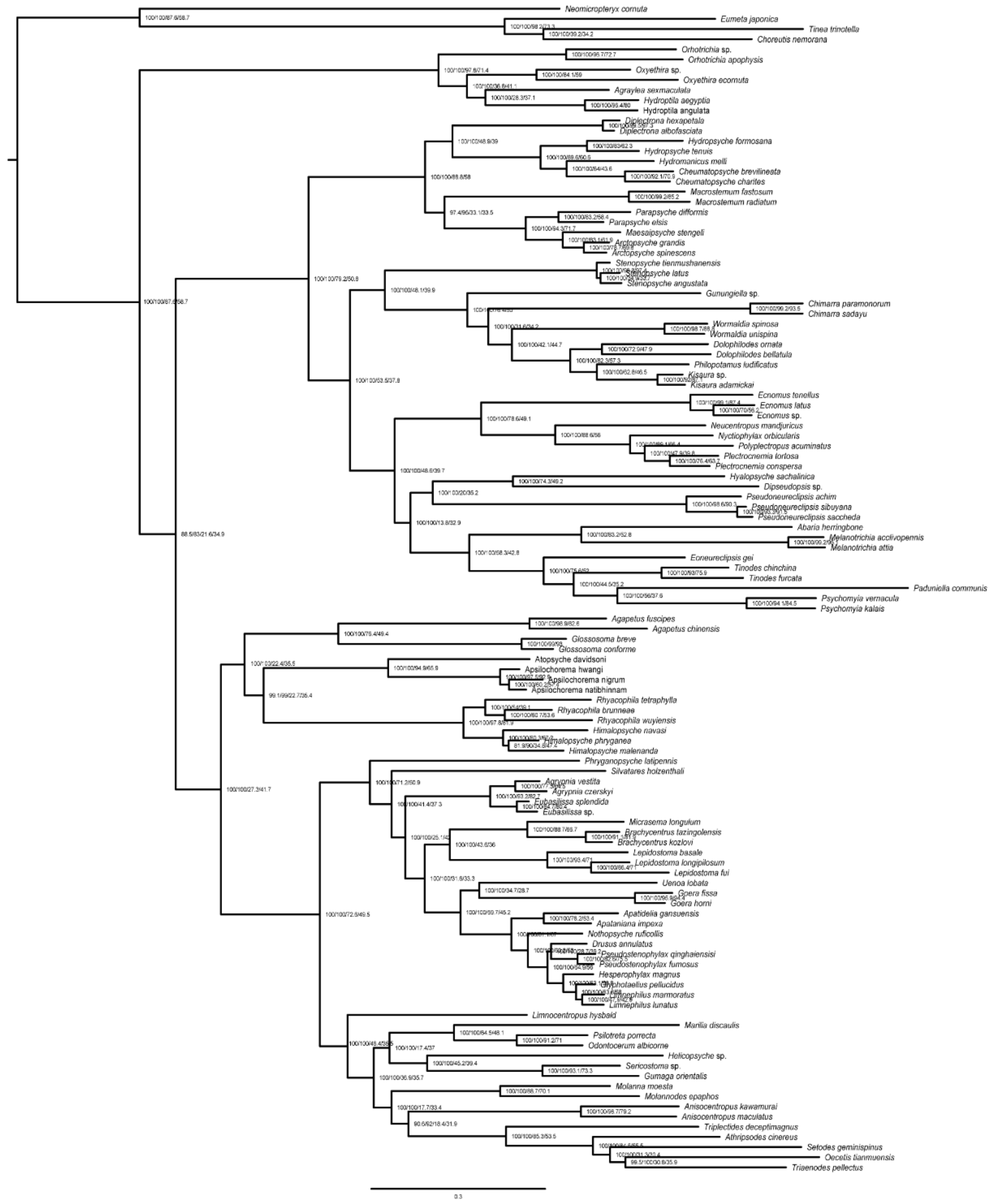

**Fig. S13** ML phylogenomic tree of Trichoptera based on USCO70_abs75 dataset with Partitioning model in IQ-TREE. Node values represent SH-aLRT/UFBoot2/gCF/sCF, respectively.

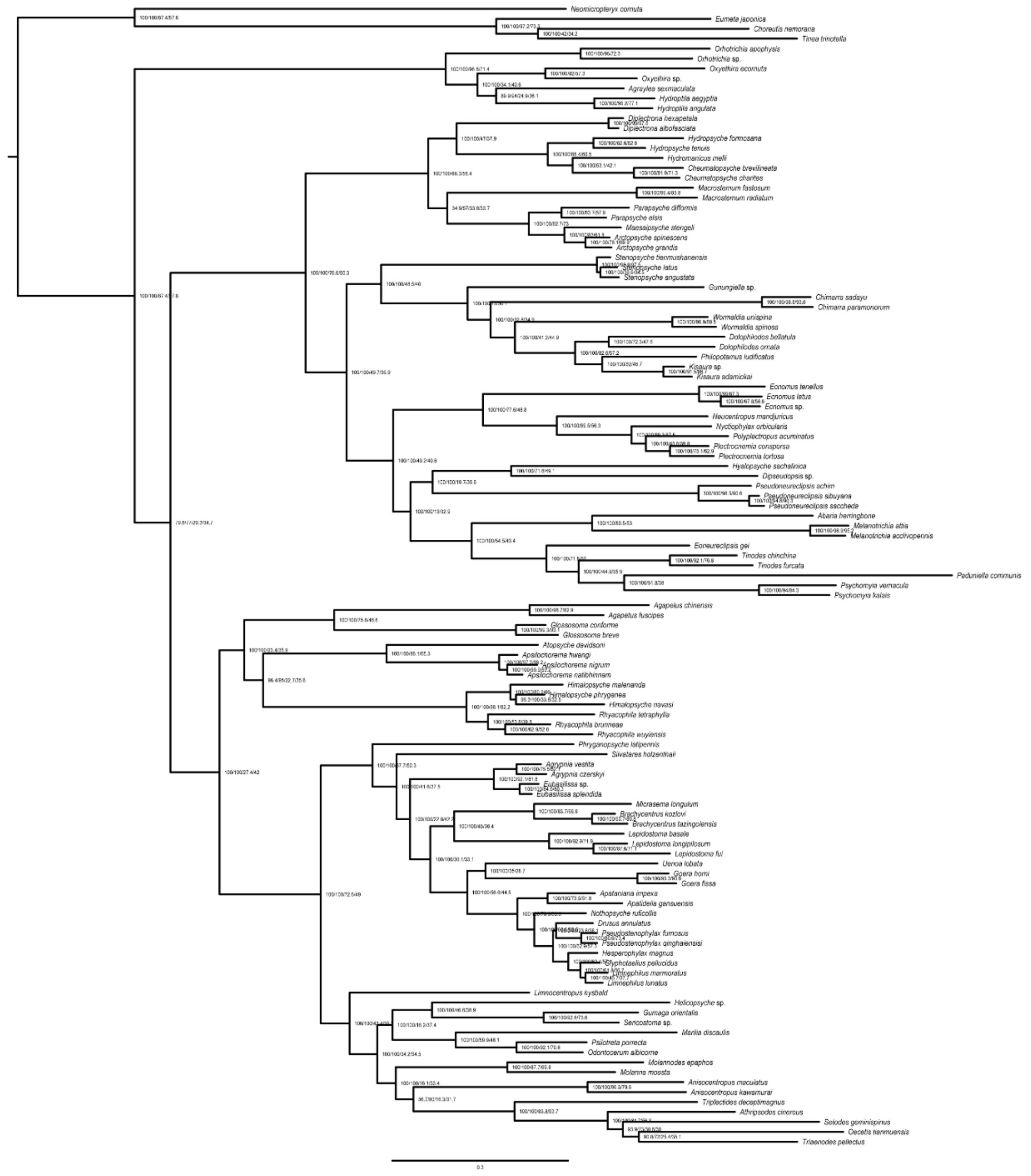


**Fig. S14** ML phylogenomic tree of Trichoptera based on USCO80_abs75 dataset with Partitioning model in IQ-TREE. Node values represent SH-aLRT/UFBoot2/gCF/sCF, respectively.

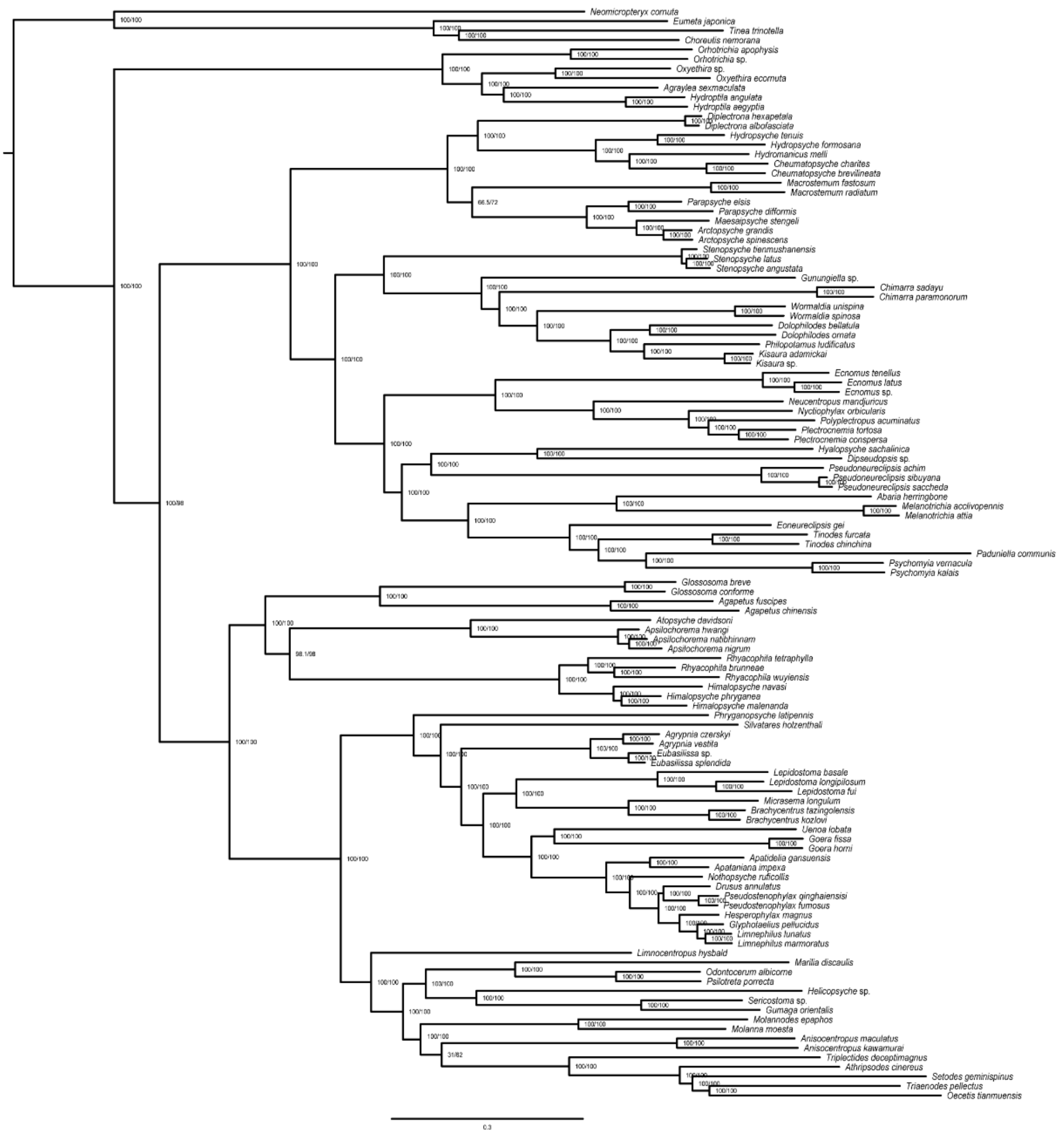


**Fig. S15** ML phylogenomic tree of Trichoptera based on USCO60_abs75 dataset with Ghost model in IQ-TREE. Node values represent SH-aLRT/UFBoot2, respectively.

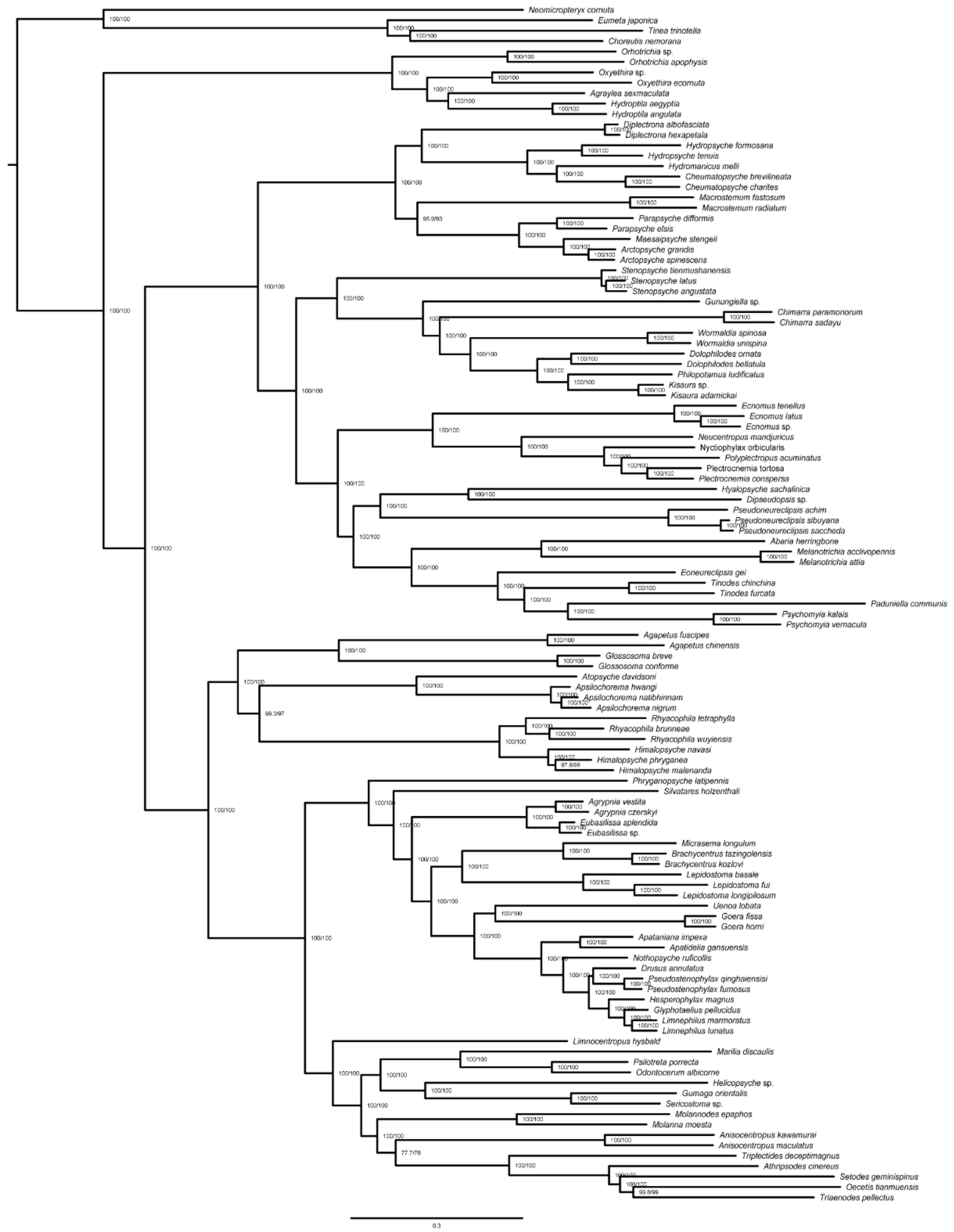

**Fig. S16** ML phylogenomic tree of Trichoptera based on USCO70_abs75 dataset with Ghost model in IQ-TREE. Node values represent SH-aLRT/UFBoot2, respectively.

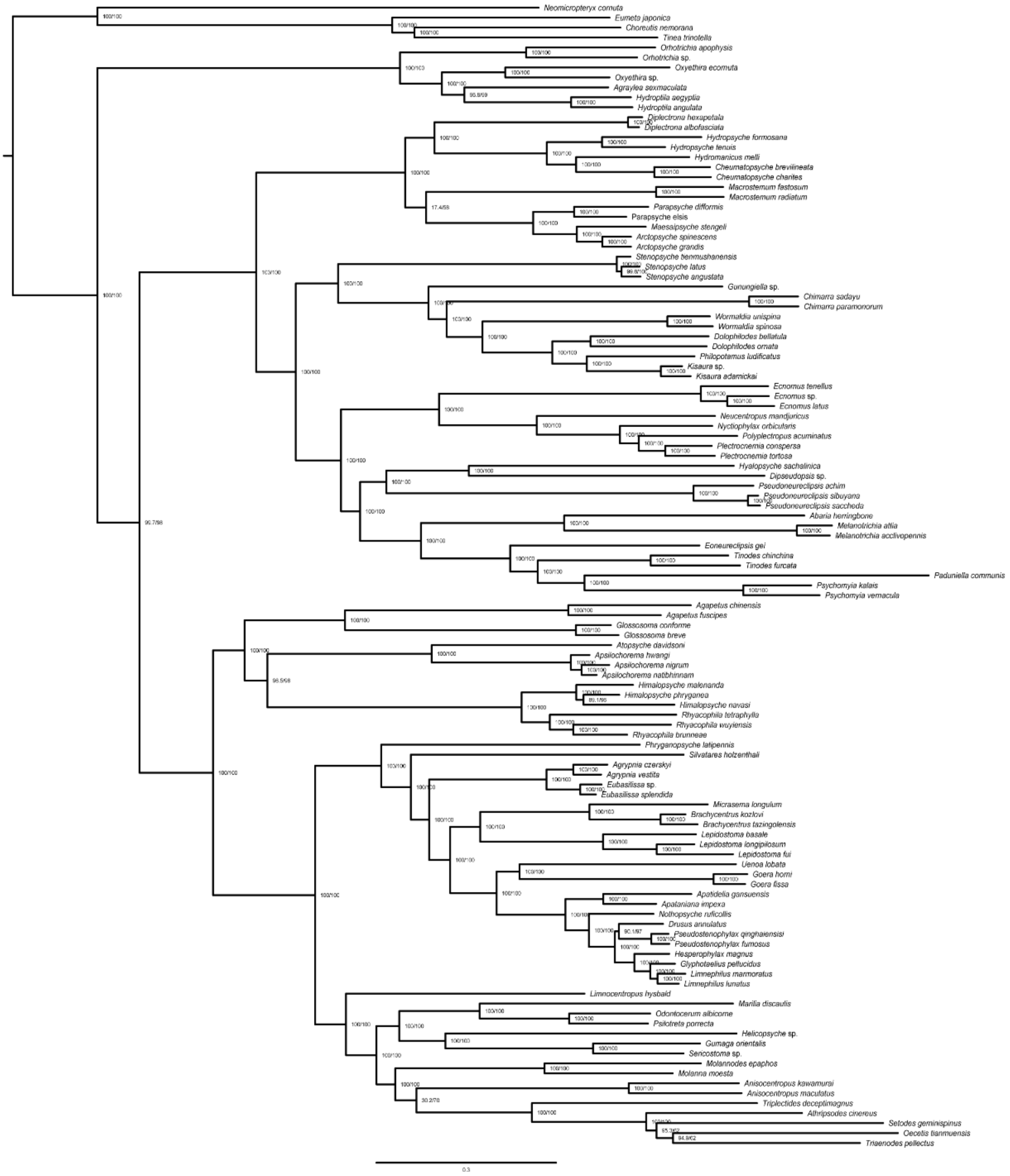


**Fig. S17** ML phylogenomic tree of Trichoptera based on USCO80_abs75 dataset with Ghost model in IQ-TREE. Node values represent SH-aLRT/UFBoot2, respectively.

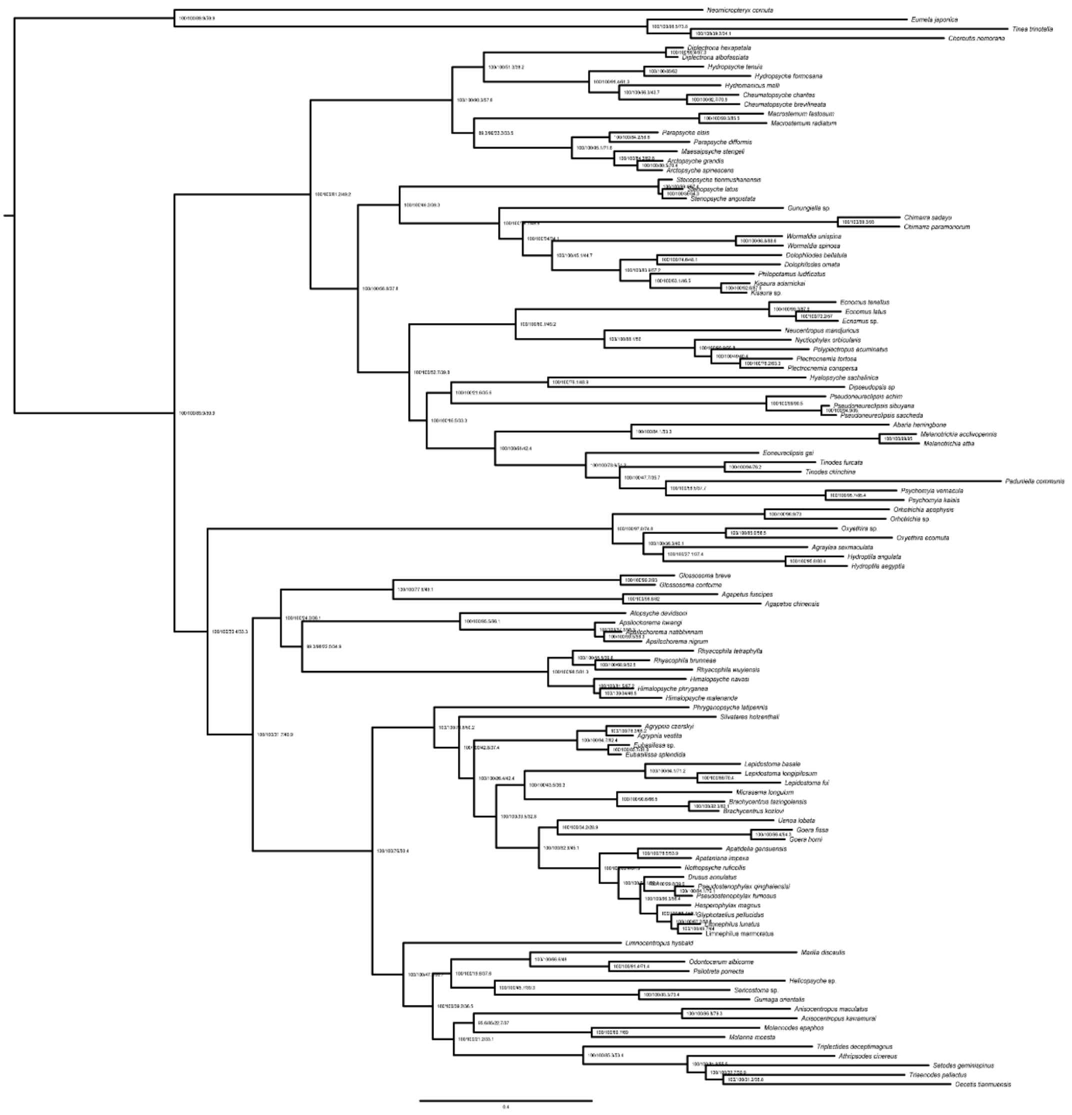

**Fig. S18** ML phylogenomic tree of Trichoptera based on USCO60_abs75 dataset with EX_EHO mix model in IQ-TREE. Node values represent SH-aLRT/UFBoot2/gCF/sCF, respectively.

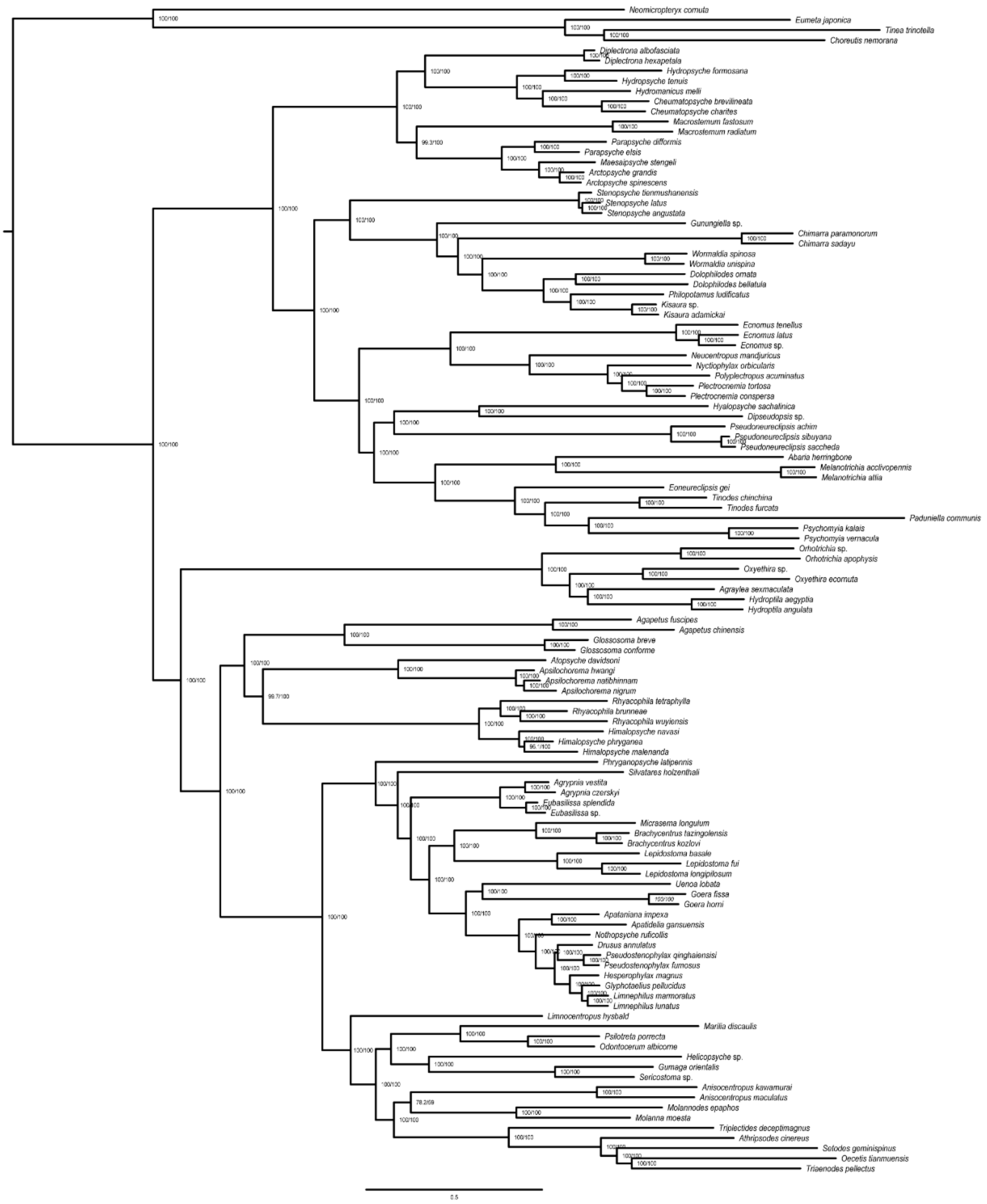


**Fig. S19** ML phylogenomic tree of Trichoptera based on USCO70_abs75 dataset with EX_EHO mix model in IQ-TREE. Node values represent SH-aLRT/UFBoot2, respectively.

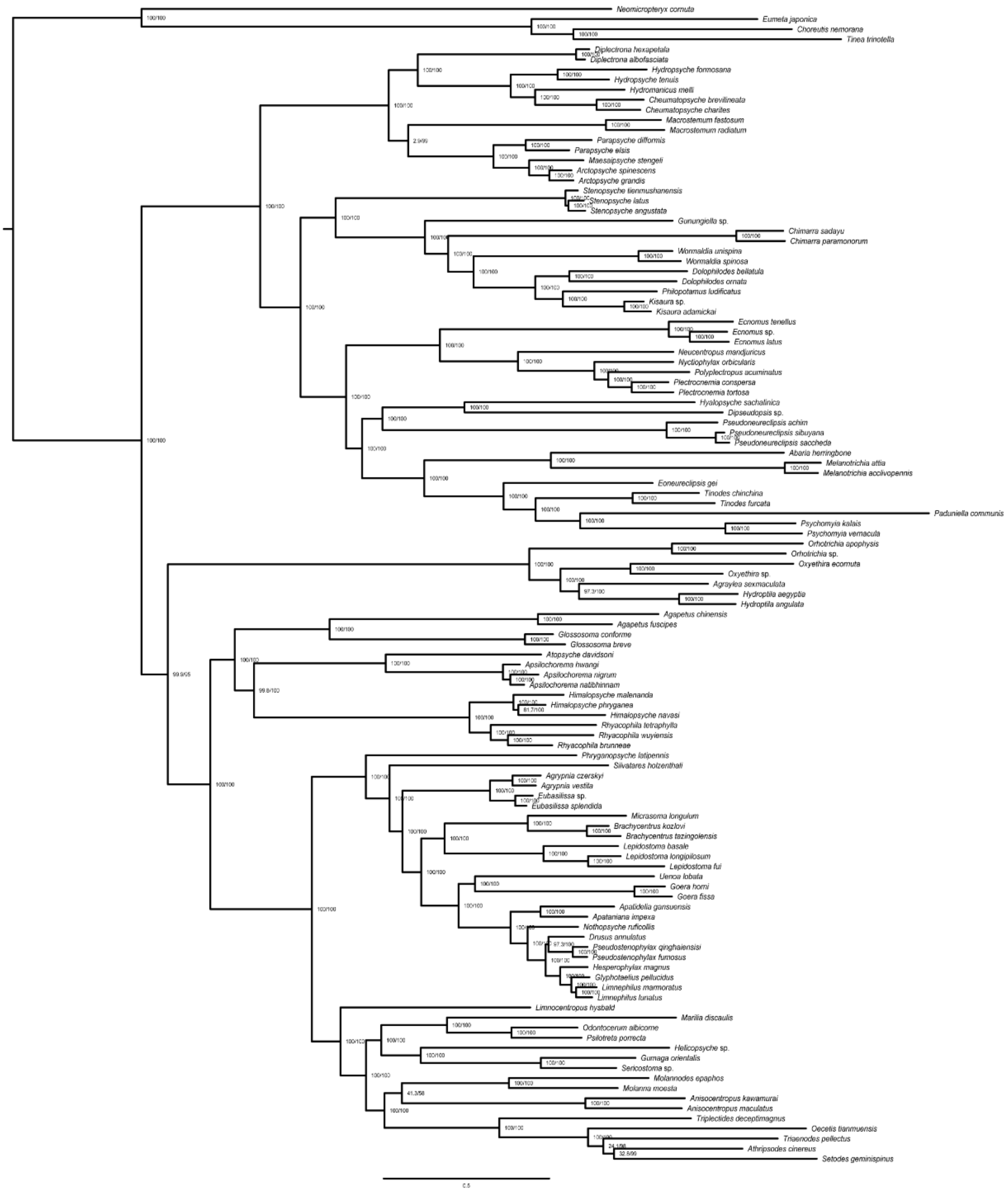


**Fig. S20** ML phylogenomic tree of Trichoptera based on USCO80_abs75 dataset with EX_EHO mix model in IQ-TREE. Node values represent SH-aLRT/UFBoot2, respectively.

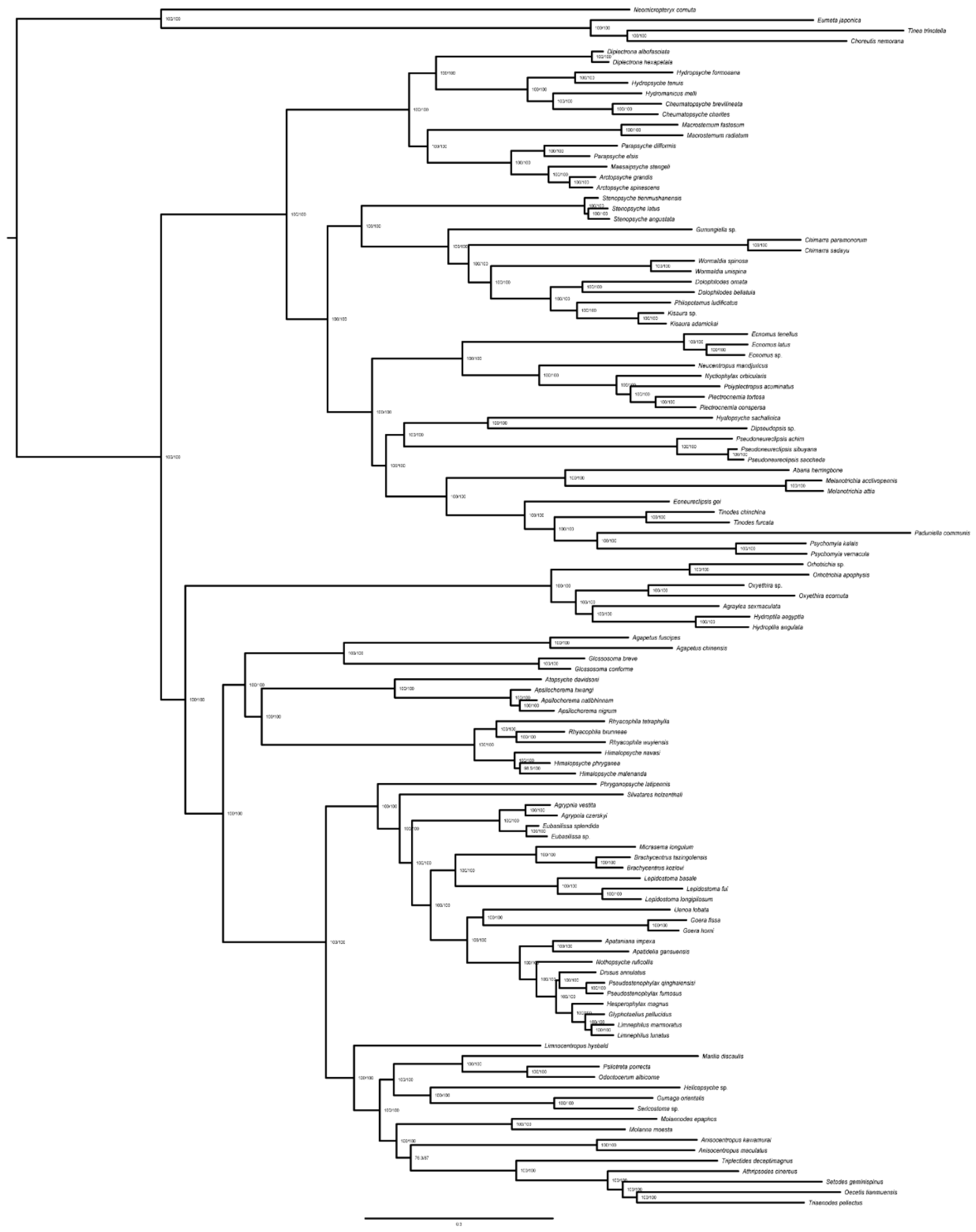

**Fig. S21** ML phylogenomic tree of Trichoptera based on USCO60_abs75 dataset with PMSF model in IQ-TREE. Node values represent SH-aLRT/UFBoot2, respectively.

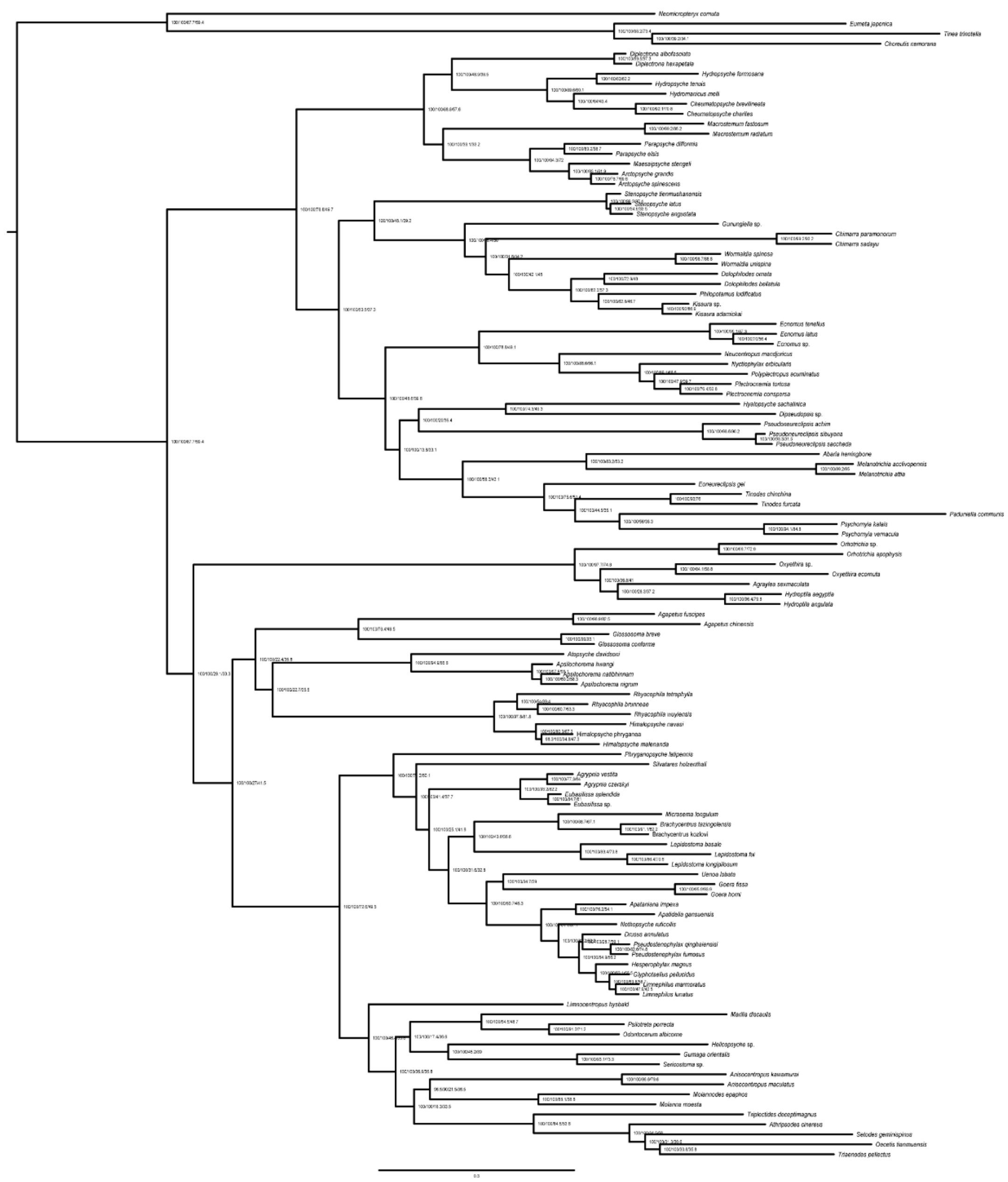

**Fig. S22** ML phylogenomic tree of Trichoptera based on USCO70_abs75 dataset with PMSF model in IQ-TREE. Node values represent SH-aLRT/UFBoot2/gCF/sCF, respectively.

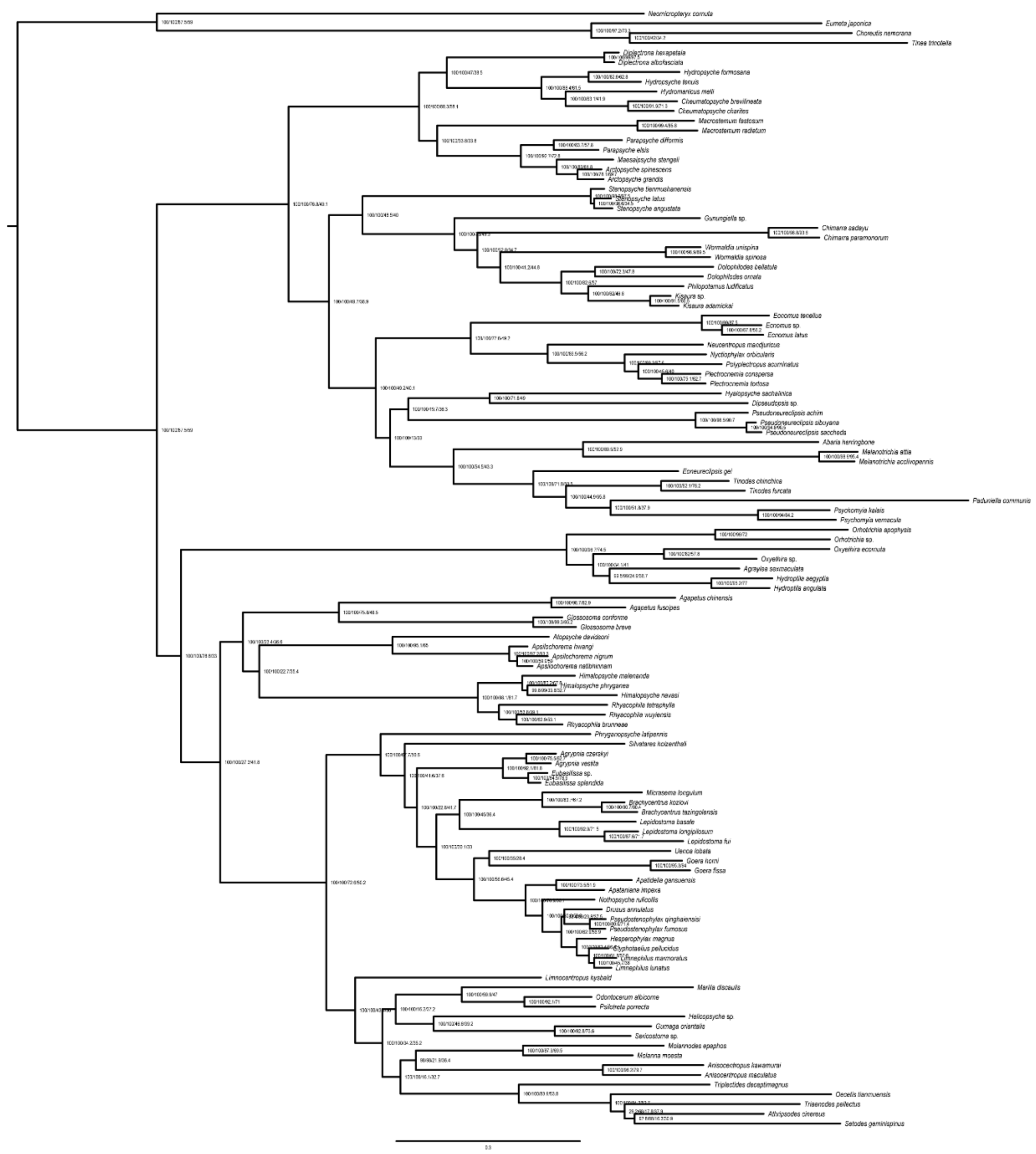

**Fig. S23** ML phylogenomic tree of Trichoptera based on USCO80_abs75 dataset with PMSF model in IQ-TREE. Node values represent SH-aLRT/UFBoot2/gCF/sCF, respectively.

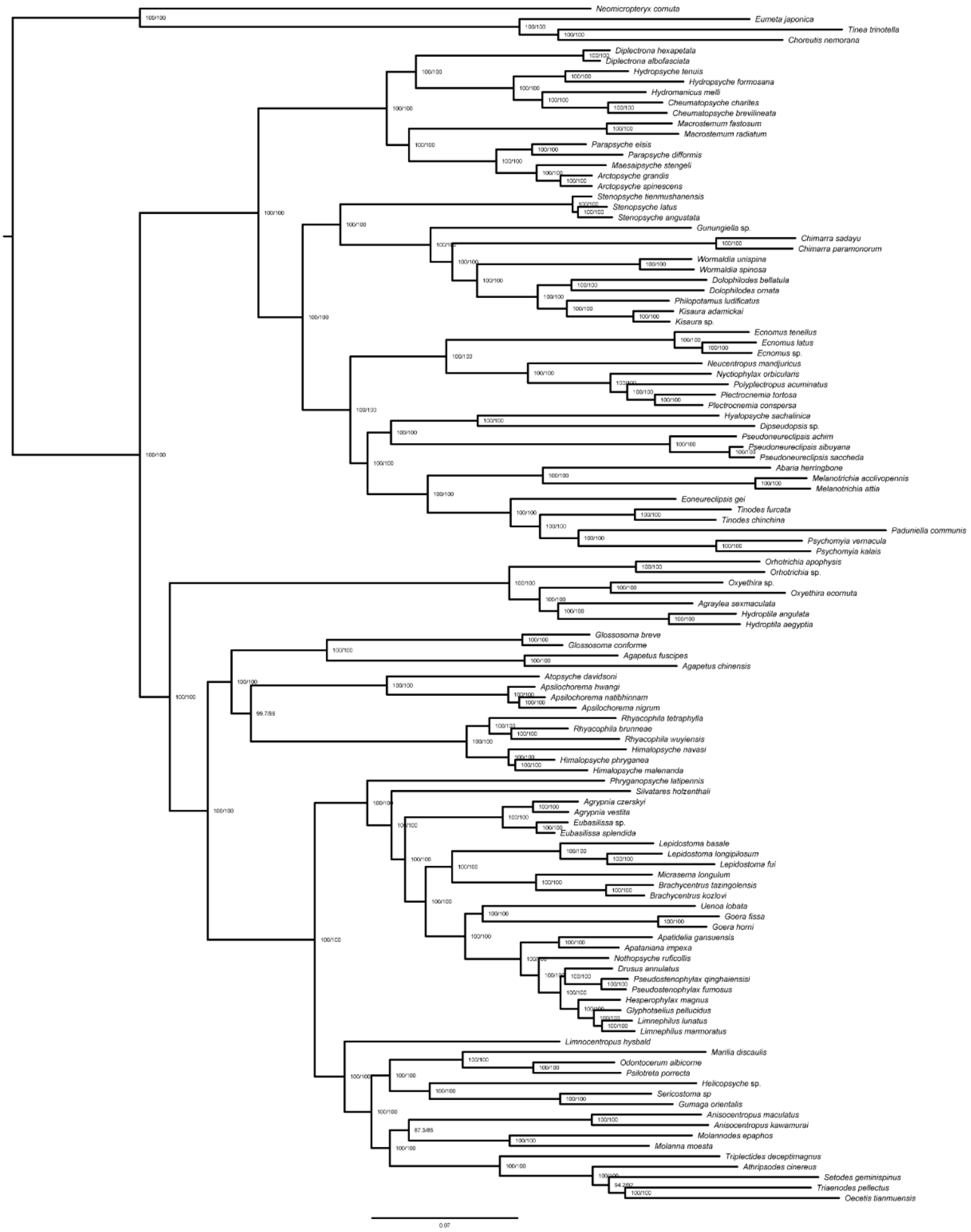

**Fig. S24** ML phylogenomic tree of Trichoptera based on USCO60_abs75 dataset with Dayhoff6 recoding model in IQ-TREE. Node values represent SH-aLRT/UFBoot2, respectively.

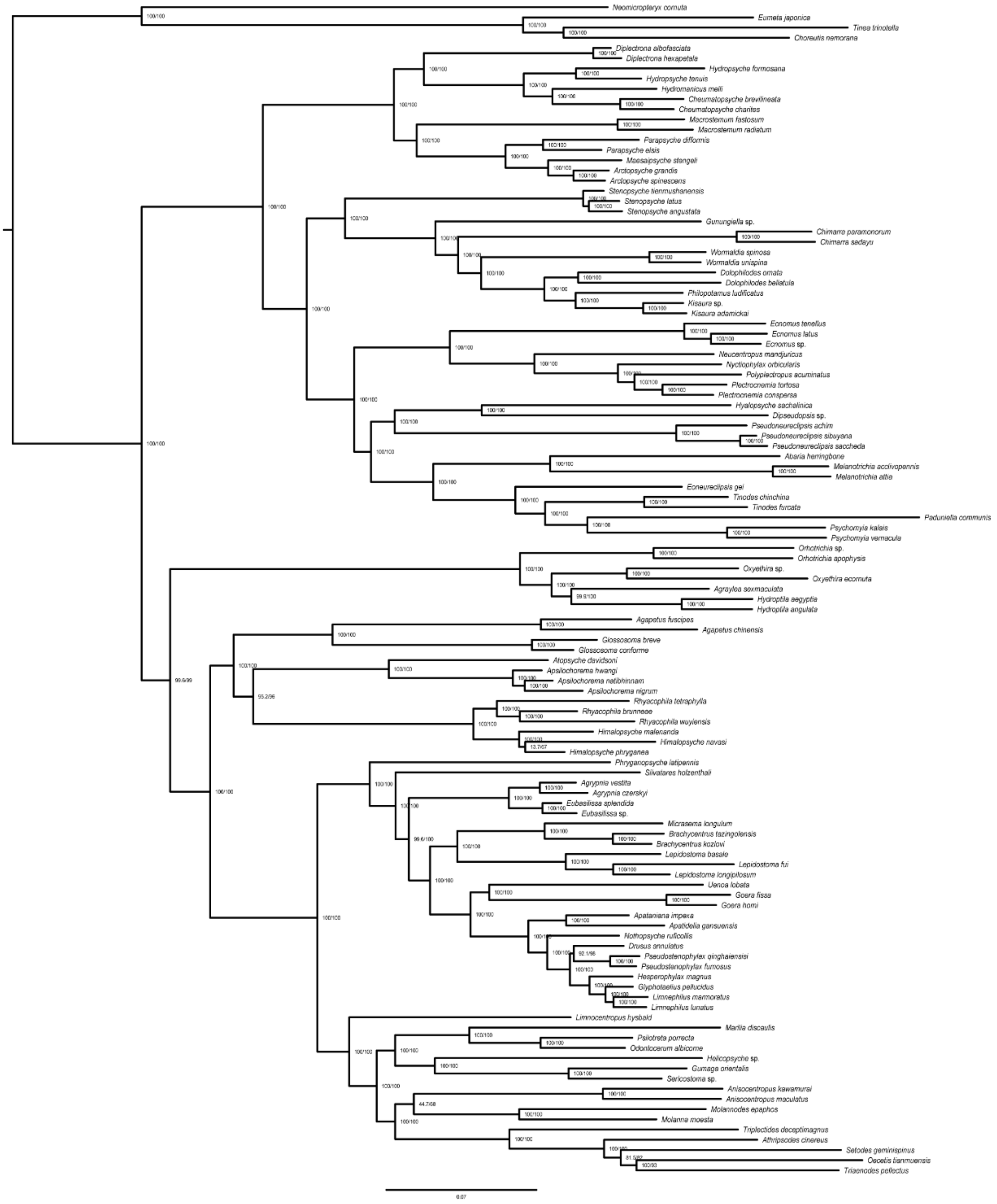


**Fig. S25** ML phylogenomic tree of Trichoptera based on USCO70_abs75 dataset with Dayhoff6 recoding model in IQ-TREE. Node values represent SH-aLRT/UFBoot2, respectively.

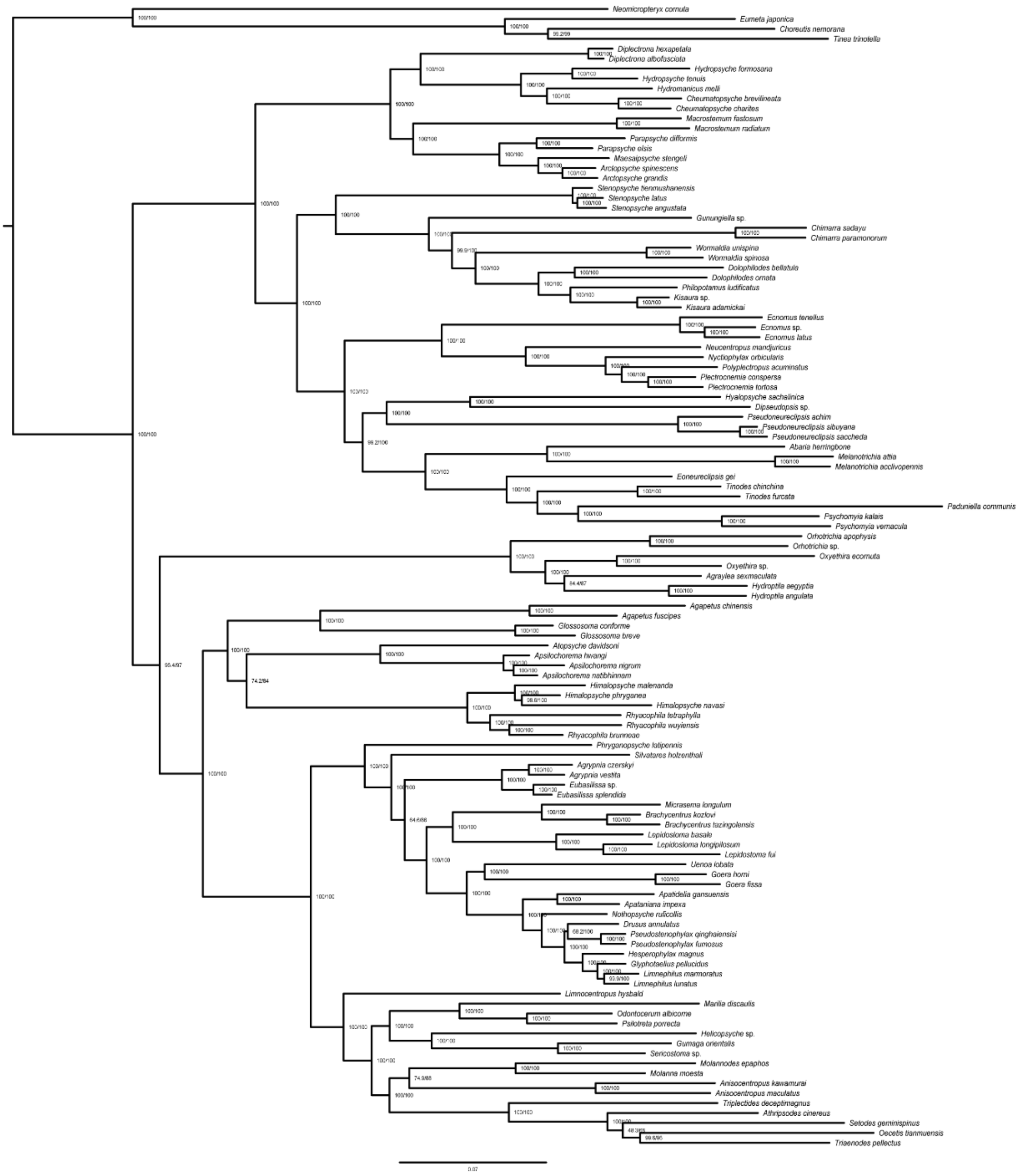


**Fig. S26** ML phylogenomic tree of Trichoptera based on USCO80_abs75 dataset with Dayhoff6 recoding model in IQ-TREE. Node values represent SH-aLRT/UFBoot2, respectively.

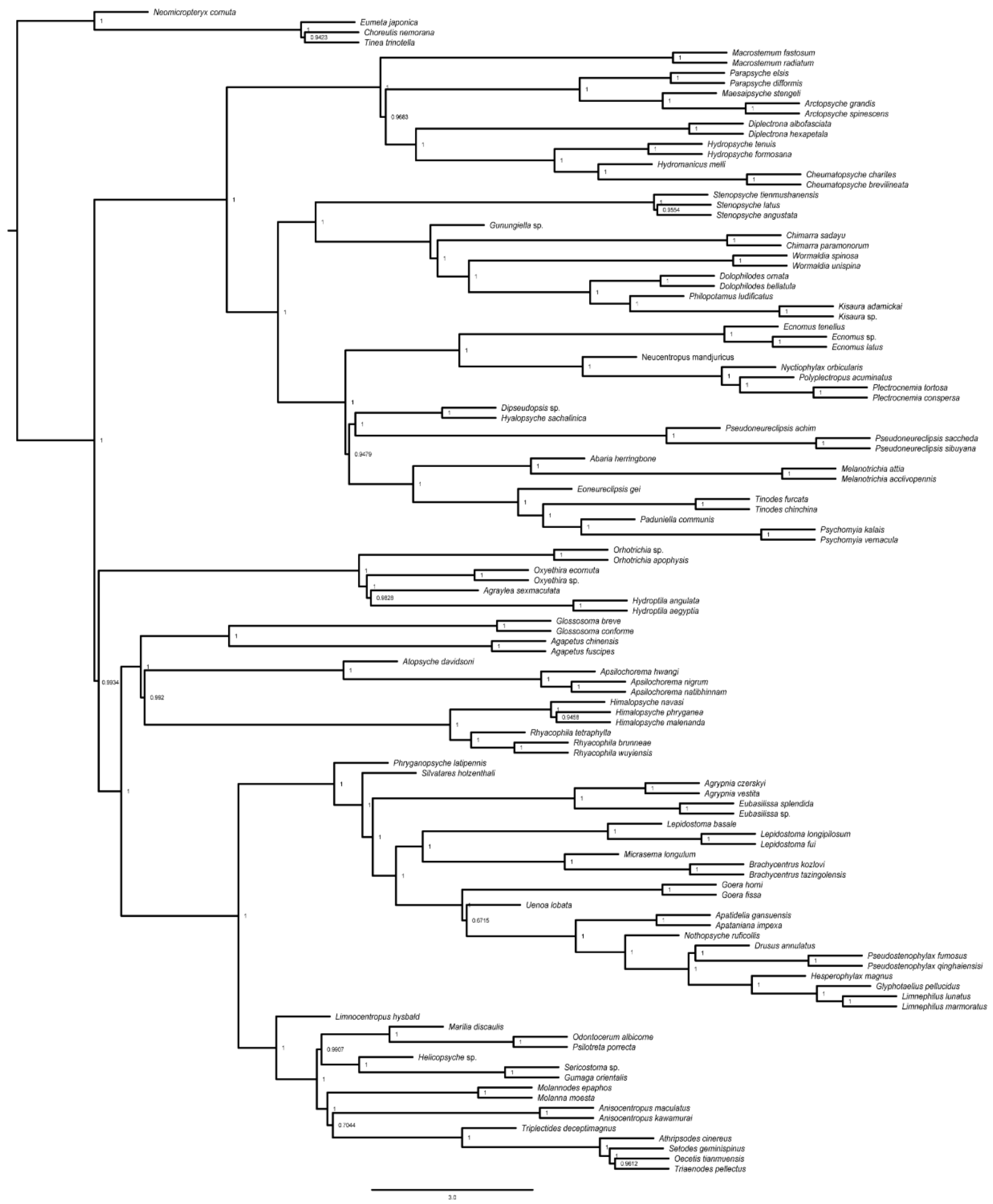

**Fig. S27** Species tree of Trichoptera based on gene tree of USCO60_abs75 using wASTRAL-hyhird. Node values represent quartet probabilities.

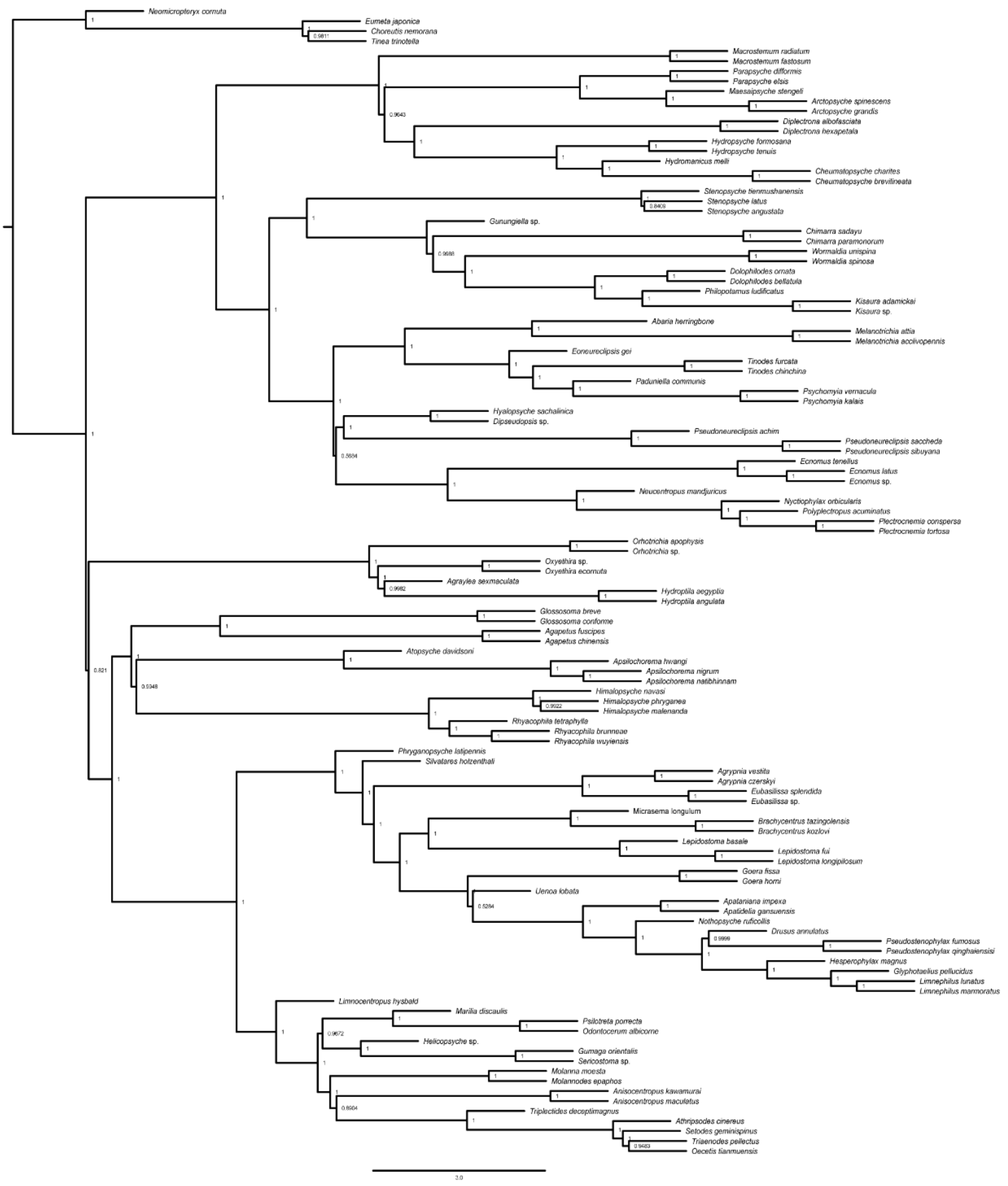


**Fig. S28** Species tree of Trichoptera based on gene tree of USCO70_abs75 using wASTRAL-hyhird. Node values represent quartet probabilities.

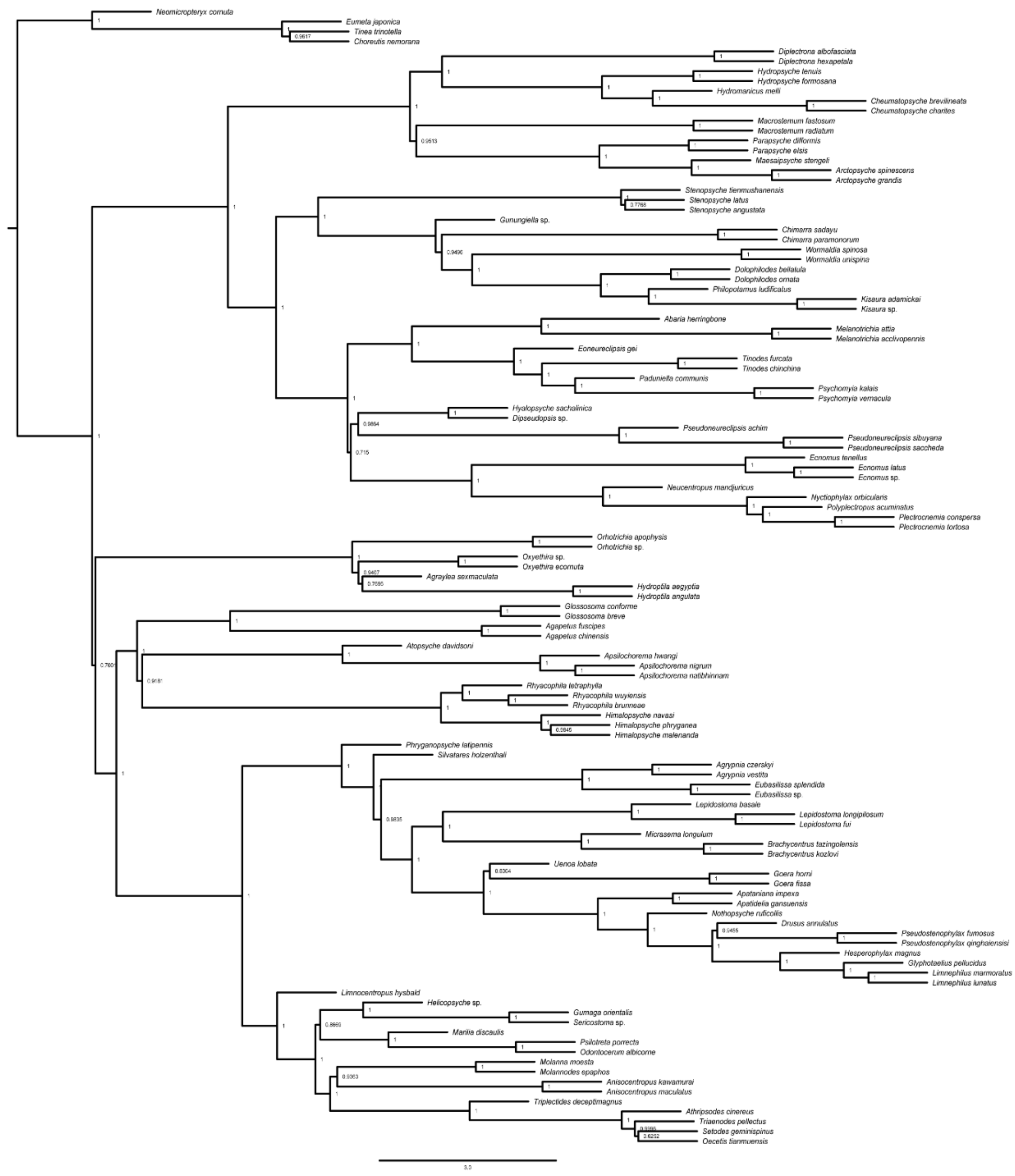

**Fig. S29** Species tree of Trichoptera based on gene tree of USCO80_abs75 using wASTRAL-hyhird. Node values represent quartet probabilities.

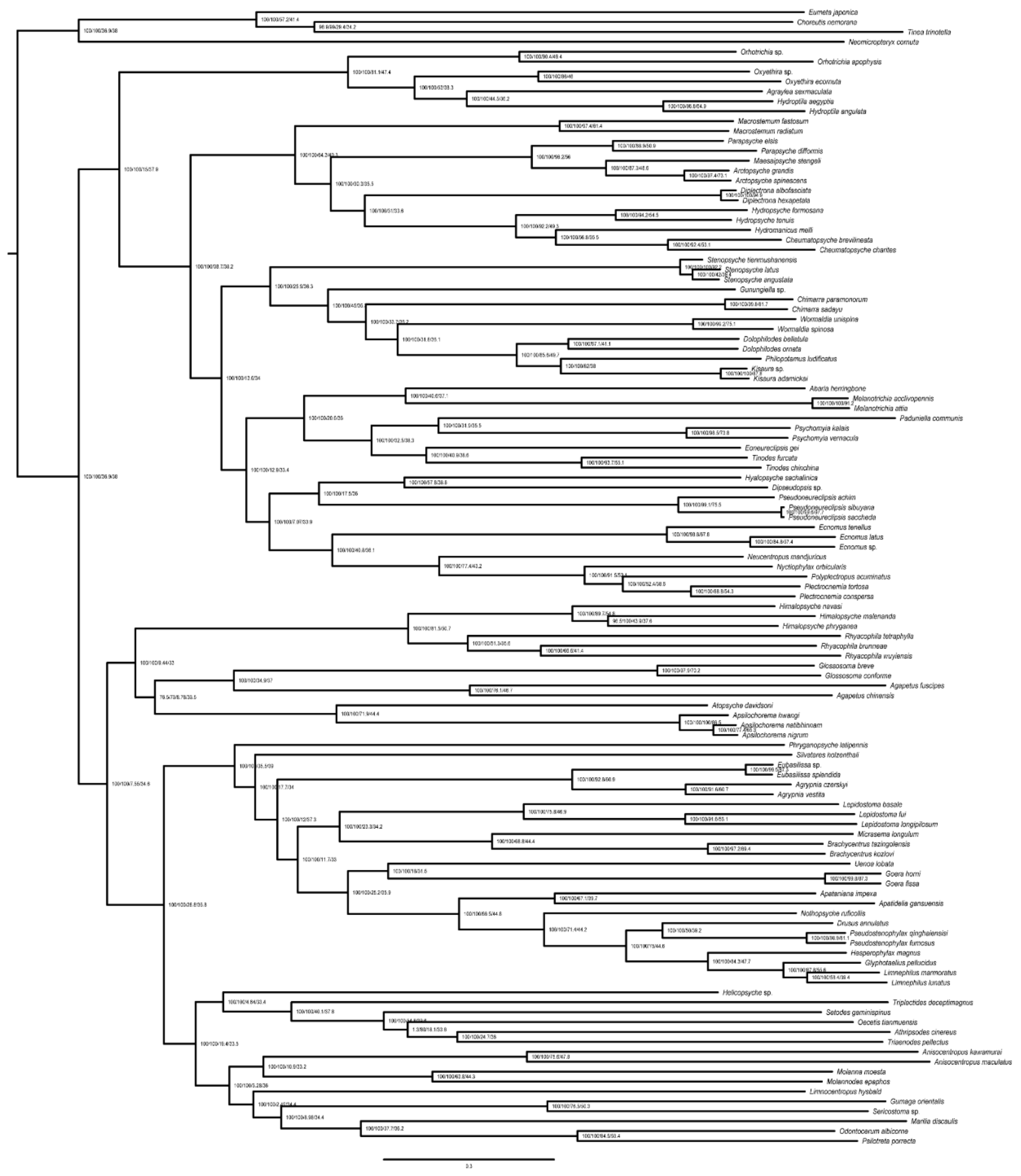

**Fig. S30** ML phylogenomic tree of Trichoptera based on UCE50_abs70 dataset with Partitioning model in IQ-TREE. Node values represent SH-aLRT/UFBoot2/gCF/sCF, respectively.

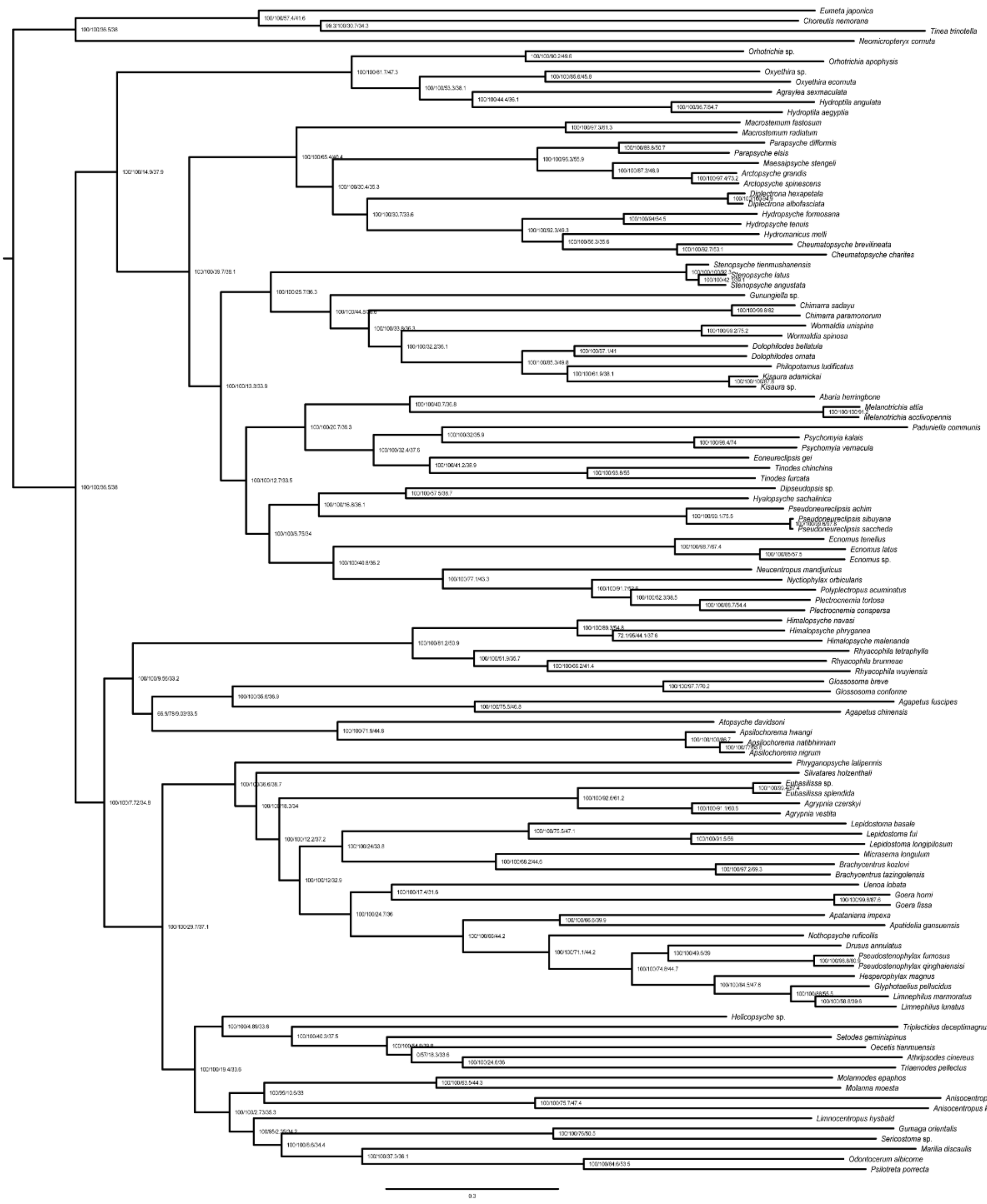


**Fig. S31** ML phylogenomic tree of Trichoptera based on UCE70_abs70 dataset with Partitioning model in IQ-TREE. Node values represent SH-aLRT/UFBoot2/gCF/sCF, respectively.

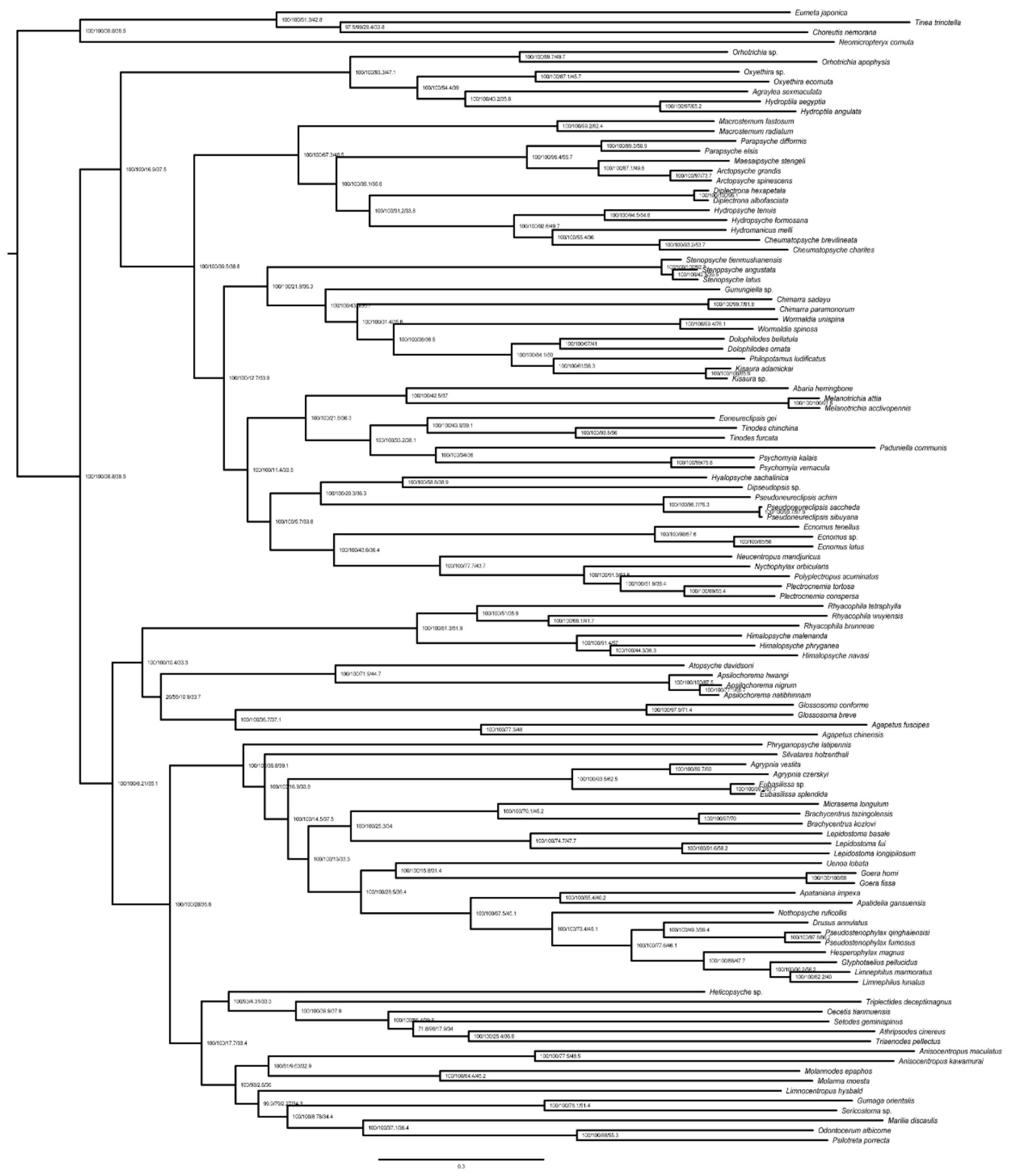

**Fig. S32** ML phylogenomic tree of Trichoptera based on UCE90_abs70 dataset with Partitioning model in IQ-TREE. Node values represent SH-aLRT/UFBoot2/gCF/sCF, respectively.

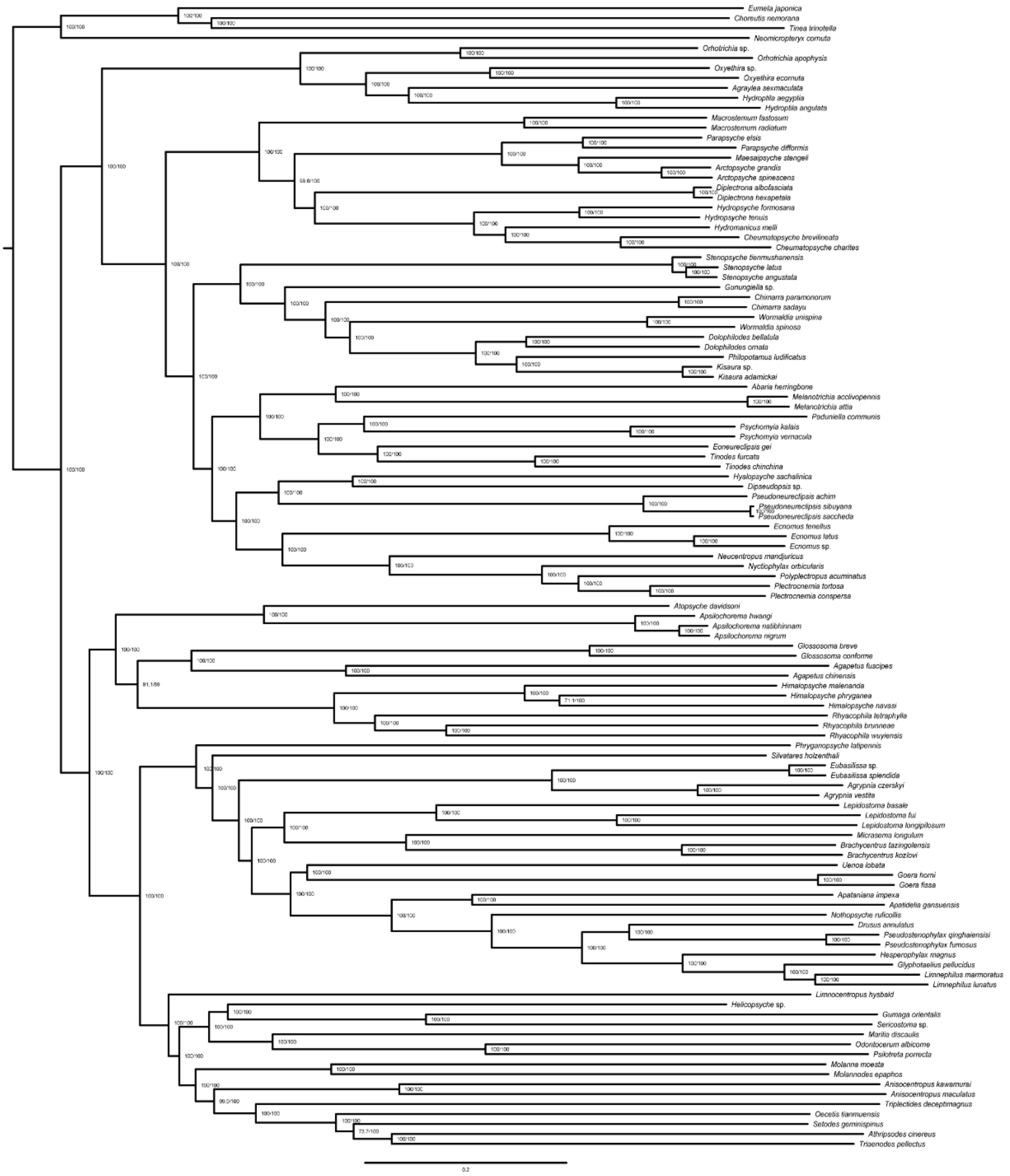


**Fig. S33** ML phylogenomic tree of Trichoptera based on UCE50_abs70 dataset with Ghost model in IQ-TREE. Node values represent SH-aLRT/UFBoot2, respectively.

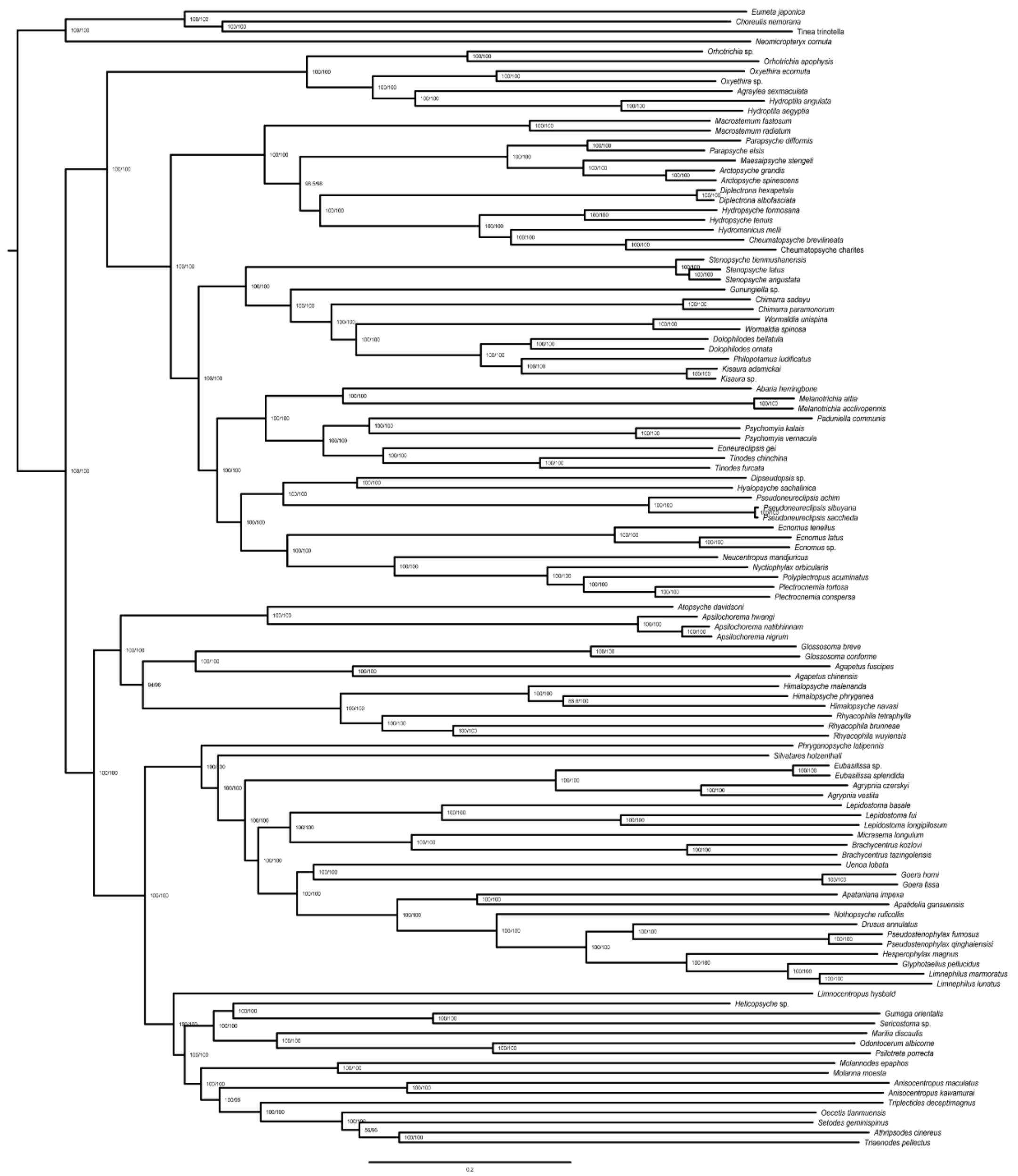


**Fig. S34** ML phylogenomic tree of Trichoptera based on UCE70_abs70 dataset with Ghost model in IQ-TREE. Node values represent SH-aLRT/UFBoot2, respectively.

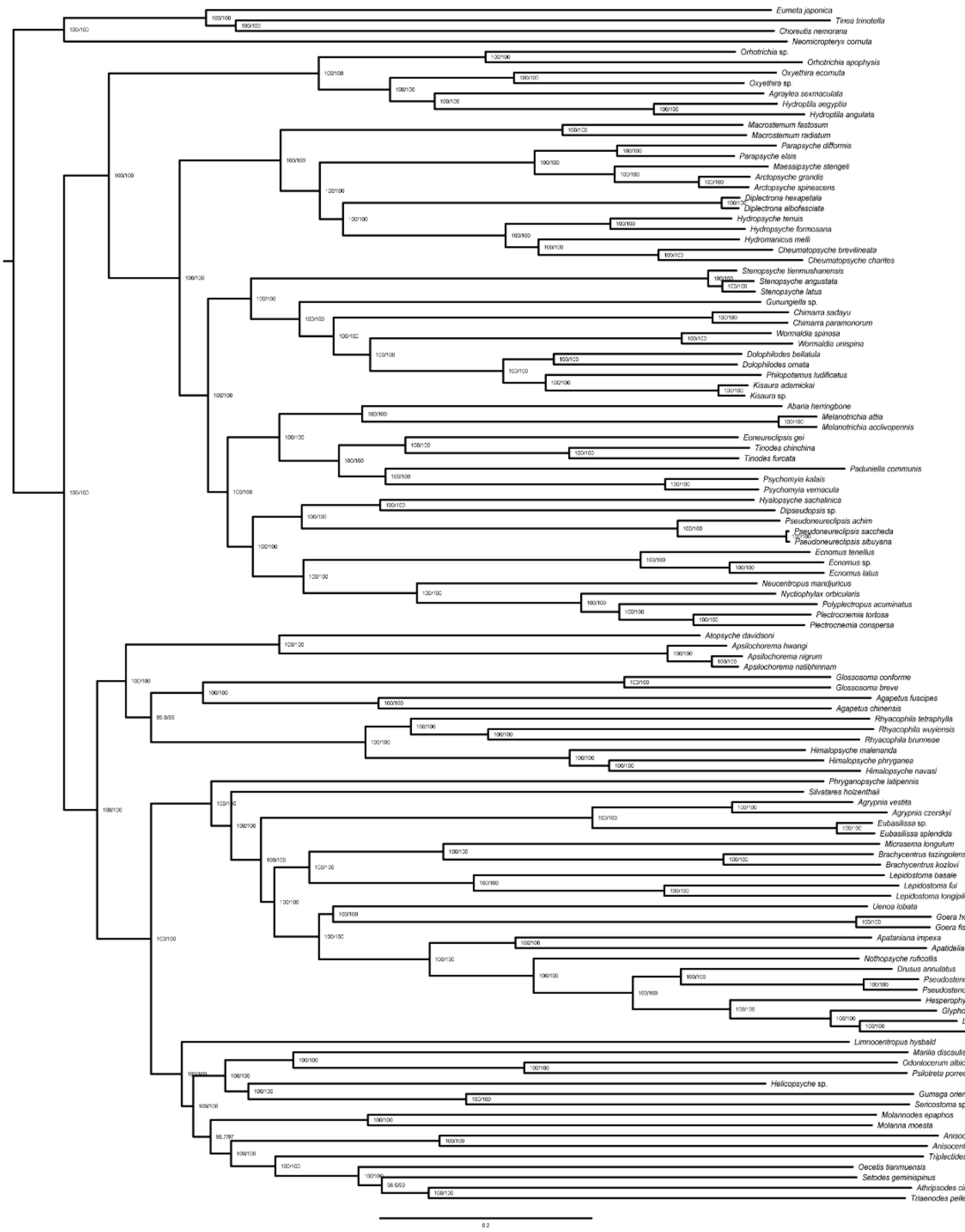


**Fig. S35** ML phylogenomic tree of Trichoptera based on UCE90_abs70 dataset with Ghost model in IQ-TREE. Node values represent SH-aLRT/UFBoot2, respectively.

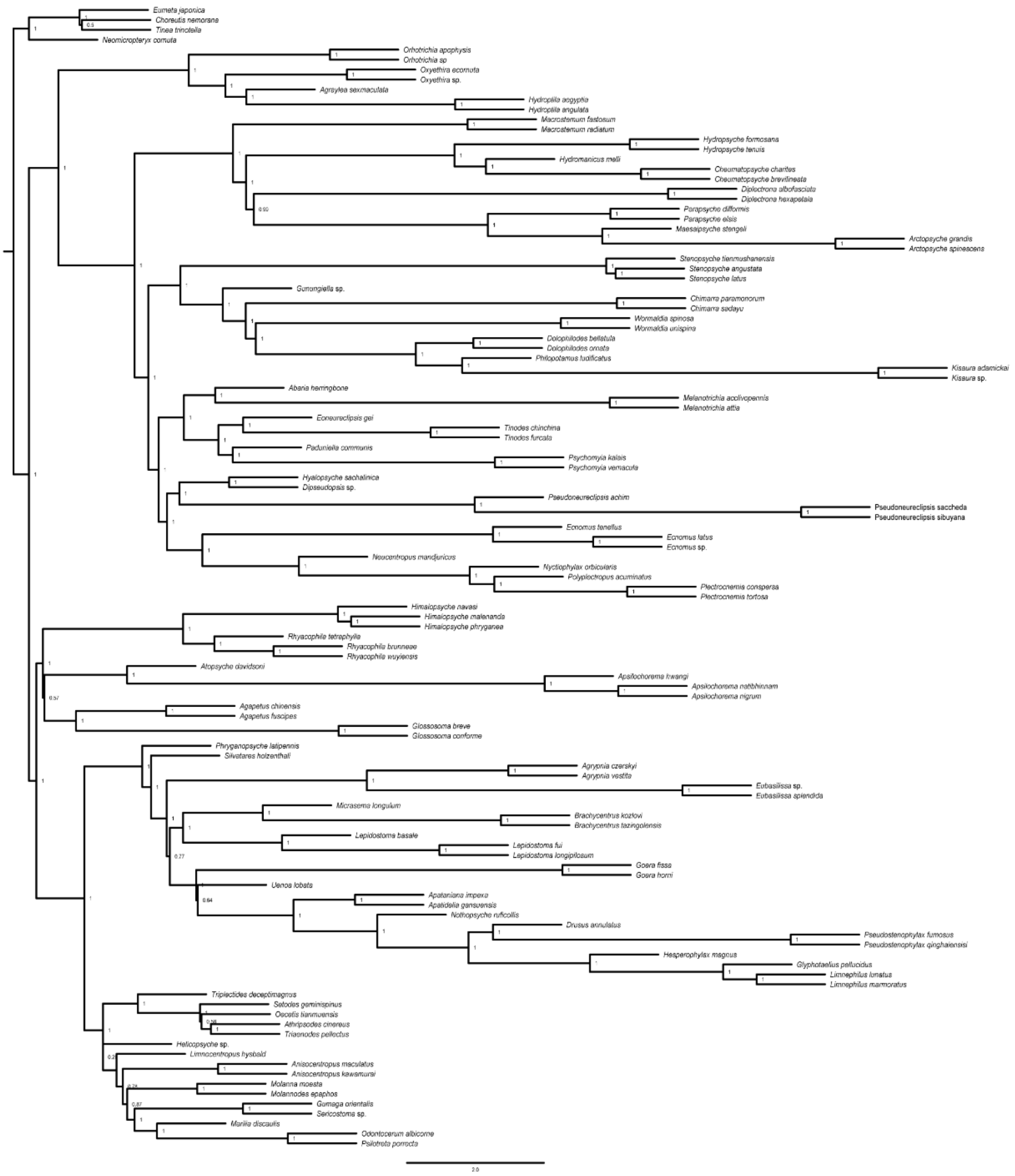


**Fig. S36** Species tree of Trichoptera based on gene tree of UCE70_abs70 using wASTRAL-hyhird. Node values represent quartet probabilities.

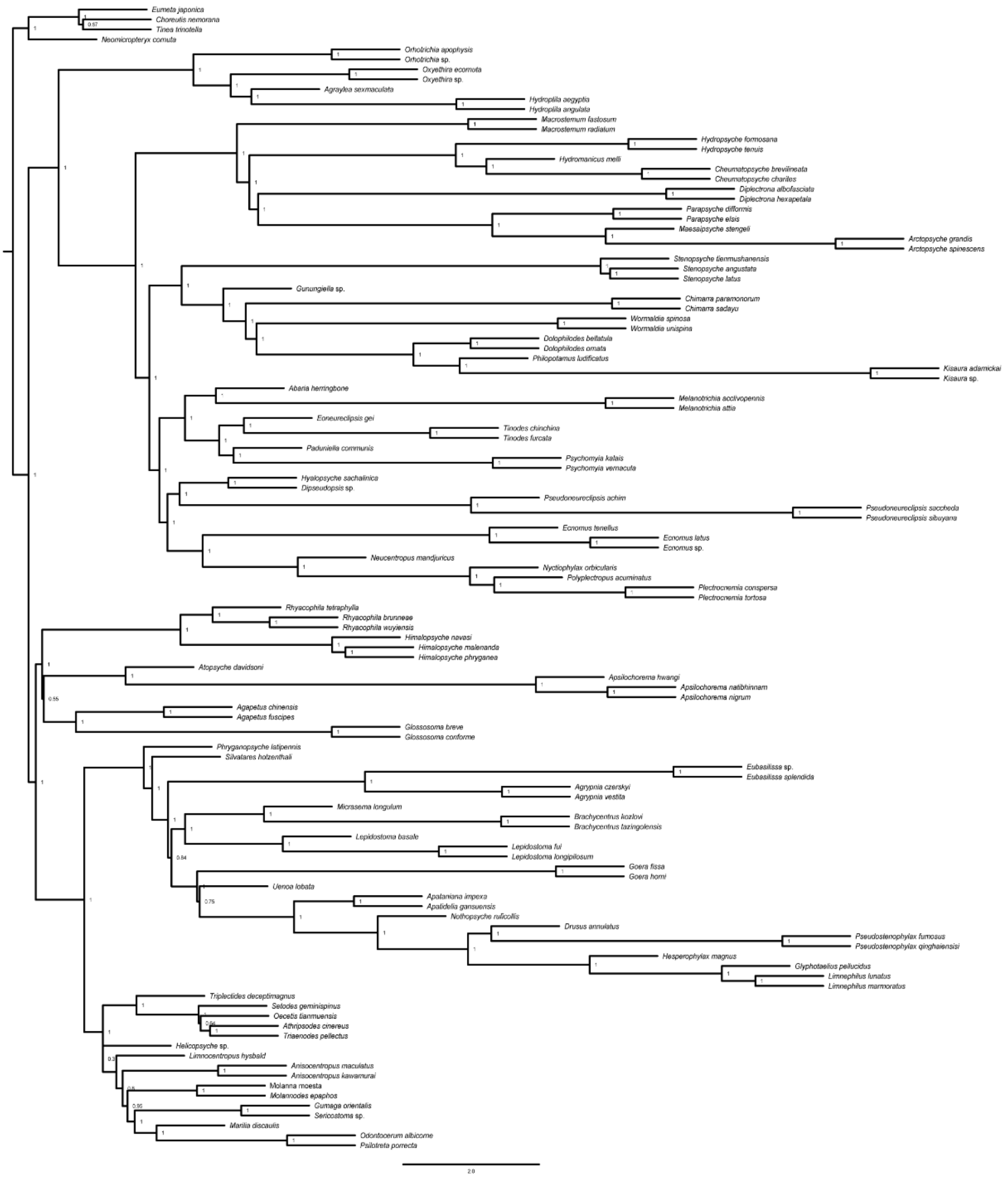


**Fig. S37** Species tree of Trichoptera based on gene tree of UCE70_abs70 using wASTRAL-hyhird. Node values represent quartet probabilities.

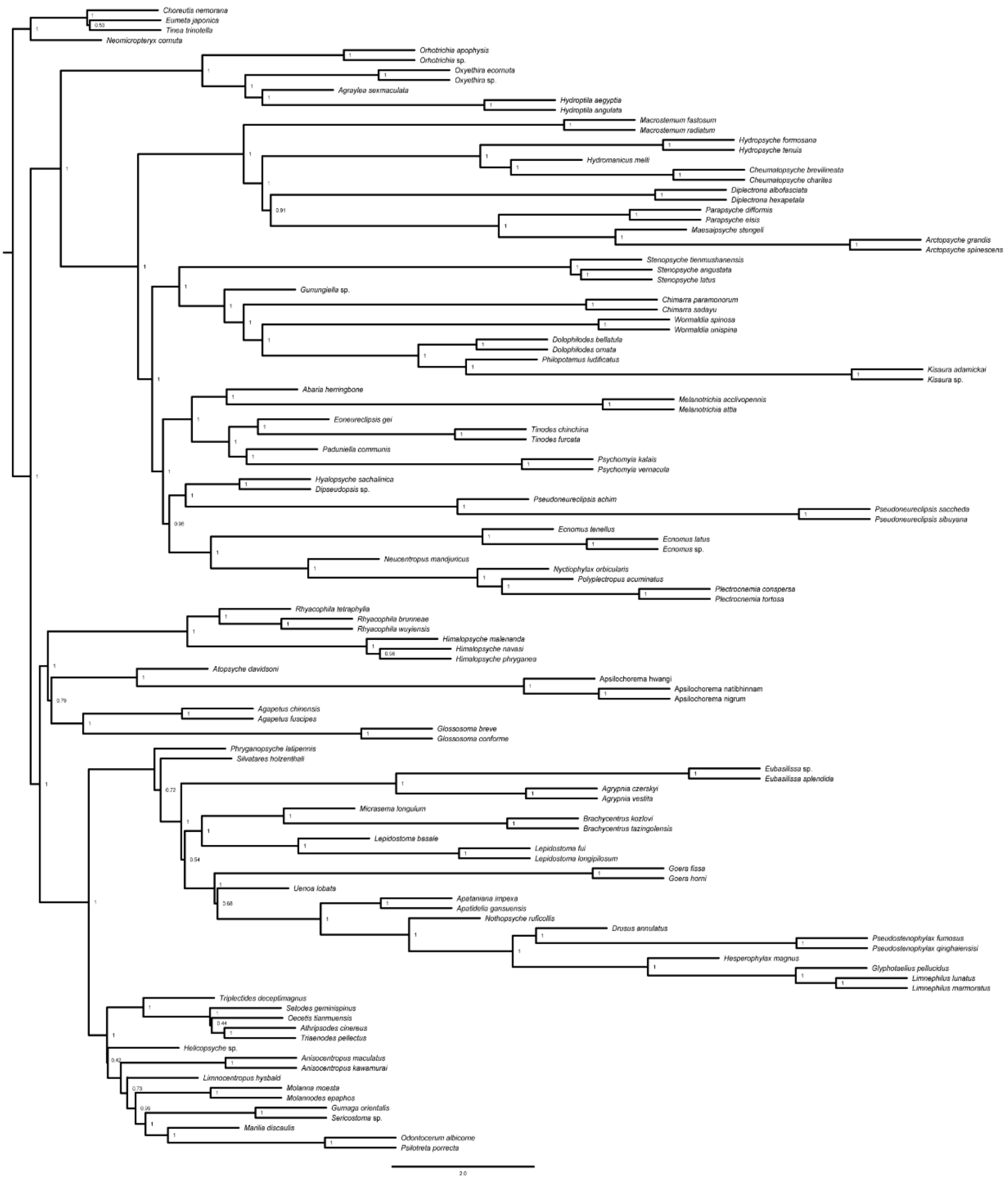

**Fig. S38** Species tree of Trichoptera based on gene tree of UCE90_abs70 using wASTRAL-hyhird. Node values represent quartet probabilities.

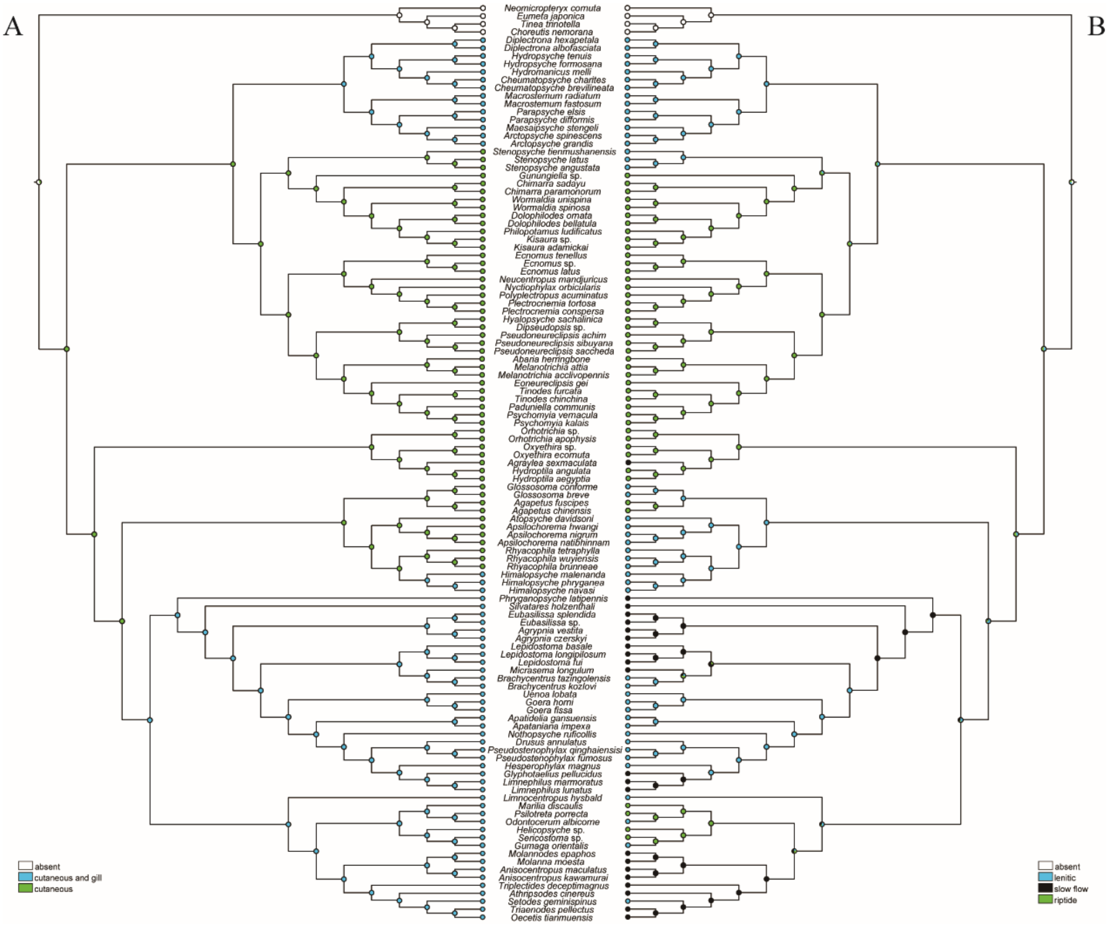


**Fig. S39** ACSR of Trichoptera. (A) Respiration; (B) Habitat. Each node indicates character states with different colorations and the proportion of the state over all examined trees.

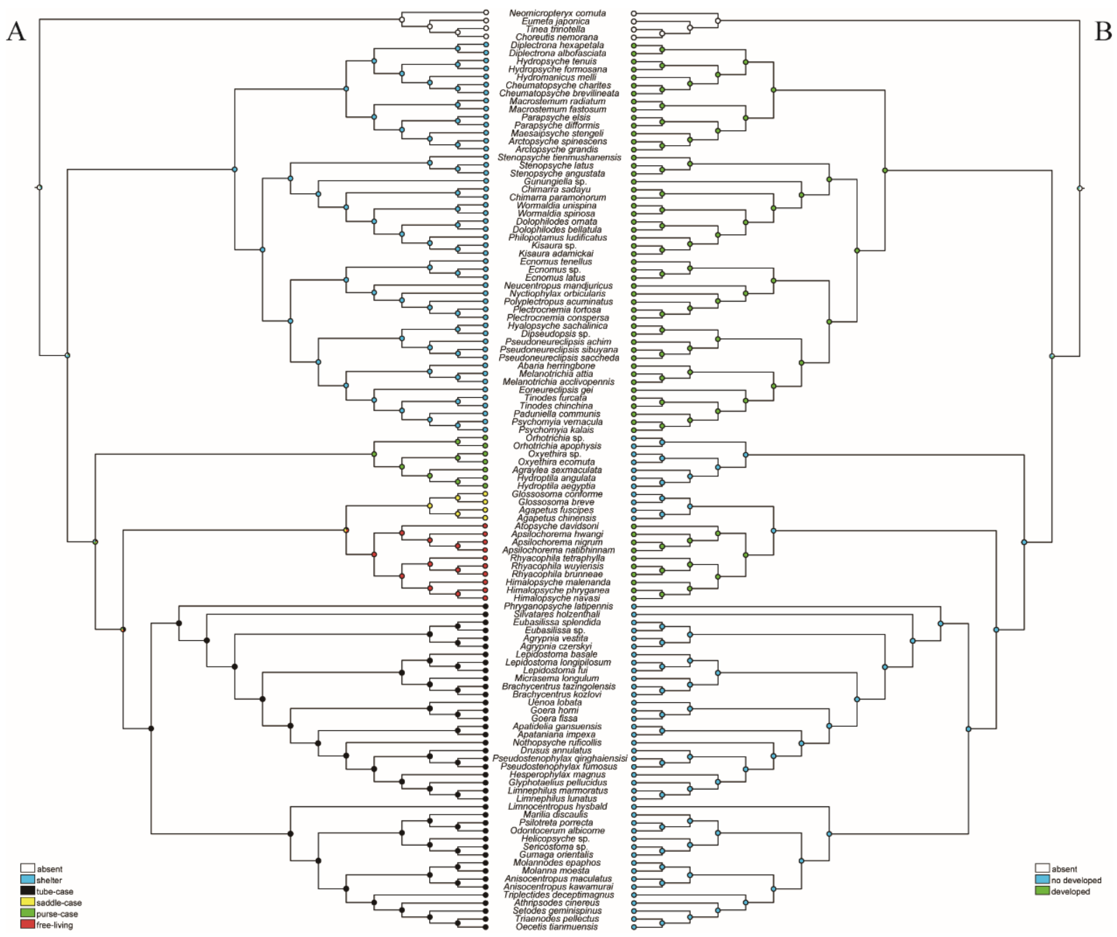


**Fig. S40** ACSR of Trichoptera. (A) Case or retreat; (B) Anal proleg. Each node indicates character states with different colorations and the proportion of the state over all examined trees.
